# Eukaryotic initiation factors eIF4F and eIF4B promote translation termination upon closed-loop formation

**DOI:** 10.1101/2024.09.10.612082

**Authors:** Ekaterina Shuvalova, Alexey Shuvalov, Walaa Al Sheikh, Alexander V. Ivanov, Nikita Biziaev, Tatiana V. Egorova, Sergey E. Dmitriev, Ilya M. Terenin, Elena Alkalaeva

## Abstract

Eukaryotic translation initiation factor eIF4F, comprising subunits eIF4G, eIF4E, and eIF4A, plays a pivotal role in the 48S preinitiation complex assembly and ribosomal scanning. Additionally, eIF4B enhances the helicase activity of eIF4A. eIF4F also interacts with PABP bound to the poly(A) tail of mRNA, thereby forming a closed-loop structure. PABP, in turn, interacts with eRF3, stimulating translation termination. Here, we employed a reconstituted mammalian system to directly demonstrate that eIF4F potently enhances translation termination. Specifically, eIF4A and eIF4B promote the loading of eRF1 into the A site of the ribosome, while eIF4G1 stimulates the GTPase activity of eRF3 and facilitates the dissociation of release factors following peptide release. We also identified MIF4G as the minimal domain required for this activity and show that eIF4G2/DAP5 can also promote termination. Our findings provide compelling evidence that the closed-loop mRNA structure facilitates translation termination, with PABP and eIF4F directly involved in this process.

## INTRODUCTION

Eukaryotic translation initiation factor 4F (eIF4F) is required for the preinitiation complex (PIC) assembly at capped mRNA and the subsequent scanning ^1–4^. eIF4F consists of eIF4G, eIF4E, and eIF4A ^5^, each of them performs a specific function in translation initiation. eIF4G is a scaffold protein which binds the other two subunits ^5^. In addition, it interacts with the eukaryotic translation initiation factor 3 (eIF3) ^6^ at the small 40S subunit of the ribosome and loads capped mRNA into 48S PIC ^7^. eIF4E recognizes the cap structure at the 5’ end of mRNA ^3,8,9^. eIF4A is an RNA helicase that unwinds the secondary structures of mRNA during scanning of the 5’ untranslated region (5’ UTR) by the PIC ^10^. The eIF4F subunits co-stimulate each other’s activity during translation initiation ^11–16^. eIF4A also associates with the eukaryotic translation initiation factor 4B (eIF4B) ^17^ which increases its helicase activity ^12,17,18^. On the other hand, the poly(A) binding protein (PABP) interacts with the N-terminal domain of eIF4G, stimulating its activity in translation initiation ^19–24^. Additionally, interaction of eIF4B and PABP was observed ^25–28^. As a result, the 5’ and 3’ ends of the mRNA get closer to each other and a closed-loop mRNA structure is formed, which has been shown to increase the translation rate ^29–32^.

In addition to the major isoform of eIF4G called eIF4G1 or eIF4GI, two other eIF4G family members, eIF4G2/DAP5 and eIF4G3/eIF4GII, have been found in mammals ^33,34^. eIF4G2/DAP5 (also known as p97 or Nat1) is involved in translation initiation along with eIF4G1. Although eIF4G2 does not bind eIF4E, it promotes cap-dependent translation via facilitating both the scanning through short upstream open reading frames (uORFs) and resumption of scanning after the translation thereof (so called translation reinitiation) ^35–37^.

Under some conditions, eIF4G1 undergoes proteolytic cleavage. For example, during infection by certain picornaviruses, e.g., poliovirus or rhinovirus, viral 2A protease cleaves eIF4G after 674 - 681 aa, which separates the N-terminal PABP- and eIF4E-binding part from the rest of the protein, thereby inhibiting cap-dependent translation ^38^. The resulting core eIF4G1 fragment p100, which contains both the MIF4G and MA3 domains, can interact with the internal ribosome entry sites (IRESs) and drive the translation of viral mRNAs. At the same time, the N-terminal fragment of eIF4G1 is believed to remain bound to the eIF4E, rendering it unresponsive to cap thus further inhibiting cap-dependent translation of cellular mRNAs ^39,40^. eIF4G1 is also cleaved by HIV protease, which cuts the protein at two sites - after 721 aa and after 1126 aa. As a result of the proteolytic cleavage three polypeptides are obtained, one of which is a p50 fragment containing the MIF4G domain, thereby also leading to inhibition of cap-dependent translation ^41–43^. Since the p100 and p50 fragments of eIF4G, as well as eIF4G2, lack the PABP binding domain, these proteins are thought to be unable to form a closed-loop mRNA structure.

PABP also interacts with eukaryotic release factor 3 (eRF3) ^44–46^. This interaction becomes possible during the formation of a termination closed-loop mRNA structure, bringing the poly(A) tail of the mRNA into close proximity with the stop codon in the ribosome ^47,48^. It was proposed that termination closed-loop structure is formed due to the structuring of the 3’ UTR, bridging the ribosome-unbound 5’ and 3’ ends of mRNA ^47^. By positioning itself near the ribosome at the stop codon, PABP stimulates translation termination ^44^.

Translation termination is performed by two eukaryotic release factors: eRF1 and eRF3. The termination process can be divided into three distinct stages: (1) recognition of the stop codons by the N domain of eRF1 ^49–53^; (2) GTP hydrolysis by eRF3 ^54–57^, which induces a conformational change in the complex of release factors and facilitates the accommodation of the M domain of eRF1 in the peptidyl transferase center (PTC) ^57^; and (3) peptidyl-tRNA hydrolysis, mediated by the GGQ loop of eRF1, with subsequent release of the newly synthesized polypeptide from the ribosome ^56^. Poly(A) tail-bound PABP has been demonstrated to stimulate translation termination by loading eRF3 or its complex with eRF1 into the ribosome ^44^.

Termination is the final stage of translation that occurs just prior to the release of the synthesized peptide from the ribosome, making its regulation critical for cellular function. Nonsense mutations can introduce stop codons within the coding sequences (CDS), resulting in what are known as premature termination codons. The latter disrupt protein synthesis and may lead to the generation of toxic truncated proteins. Additionally, ribosomes may encounter stop codons in the uORFs within 5’ UTRs or in the small downstream open reading frames (dORFs) within 3’ UTRs, as well as in extentions of the CDSs within 3’ UTRs if the main CDS stop codon experiences readthrough. These scenarios highlight the importance of precise control over the termination stage of translation. We propose that these situations require auxiliary proteins to prevent undesirable translation termination or to support an appropriate peptide release. In addition to PABP, several other proteins have been implicated in translation termination, including the yeast ribosome recycling factor Rli1 ^58^, the human and yeast nuclear mRNA export factors DDX19 and Dbp5 ^59,60^, the human and yeast translation initiation factors eIF3j and Hcr1 ^61,62^, the human translation factor eIF5A ^63^, the human programmed cell death 4 protein (PDCD4) ^64^, and the SARS-CoV-2 viral protein Nsp1 ^65^. Notably, the PABP-binding proteins PAIP1 and PAIP2 have emerged as critical regulators in inhibiting termination at premature termination codons ^66^.

However, the regulation of translation termination across various parts of mRNA and under different cellular conditions (such as apoptosis, cellular stress, virus infection, etc.) remains inadequately explored. A closed-loop structure formation can bring the initiation and termination ribosomal complexes close to each other ^47,48^. This raises the question: could initiation factors that interact with PABP also play a role in the regulation of translation termination?

To address this question, we examined the activities of release factors in the presence of the human eIF4F complex and its individual subunits. Using a reconstituted mammalian translation system and pretermination complexes isolated from rabbit reticulocyte lysate (RRL), we show that eIF4G1, eIF4E, eIF4A, eIF4B, as well as the eIF4G1 and eIF4G2 deletion variants (p100, p50 and p86) affect various stages of translation termination. Furthermore, we investigated the ability of initiation factors to bind release factors and pretermination or termination ribosomal complexes. Based on these findings, we propose a model for the regulation of translation termination by eIF4F and eIF4B, highlighting their potential roles in coordinating the termination process.

## RESULTS

### eIF4F promotes termination of translation

To investigate activity of eIF4F in translation termination, we co-expressed all three subunits of human eIF4F (eIF4G1, eIF4A, and eIF4E) in a baculovirus expression system and purified the trimer by affinity and ion exchange chromatographies. The eIF4F preparation contained all the subunits and was free of human release factors eRF1 and eRF3a (Fig. S1A). The functional activity in translation initiation of purified eIF4F was confirmed by the assembly of the 48S initiation complex in a reconstituted translation system (Fig. S2).

First of all, we studied if eIF4F affects peptidyl-tRNA hydrolysis induced by release factors. For this purpose, we employed the Termi-Luc assay, which allowed us to analyze the efficiency of release of the nanoluciferase (Nluc) during termination of translation from preTC isolated from RRL ^67^. PreTCs were assembled at uncapped but polyadenylated Nluc mRNA at the UAA stop codon and then purified by sucrose density gradient ultracentrifugation. As we previously showed, such preTCs contain PABP associated with the poly(A) tail and ribosomal proteins but not initiation factors ^47^. This allowed us to study the functions of eIF4F exclusively in translation termination. eIF4F was added to Nluc preTC in the presence of either eRF1 alone or eRF1 with eRF3a-GTP (Fig. 1A,B, S1B). We have shown that eIF4F significantly (up to 3.5-fold) increased the rate of Nluc release from the ribosome in the presence of both eRF1 and eRF3a (we observed this effect even at a molar ratio of as low as 1:1 for eRFs to eIF4F) (Fig. 1A). eIF4F also stimulated translation termination in the presence of eRF1 alone, but to a smaller extent (Fig. 1B). An increase in peptide release was detected in the eRF1:eIF4F ratio of 1:2. In the absence of release factors, eIF4F did not induce peptidyl-tRNA hydrolysis on its own (Fig. S1C). Glutathione S-transferase (GST) was used as a negative control and had no effect on translation termination (Fig. S1D). Collectively, these findings indicate that human eIF4F stimulates peptide release both in the presence of eRF1 alone and in the presence of the eRF1-eRF3 complex.

**Figure 1.**
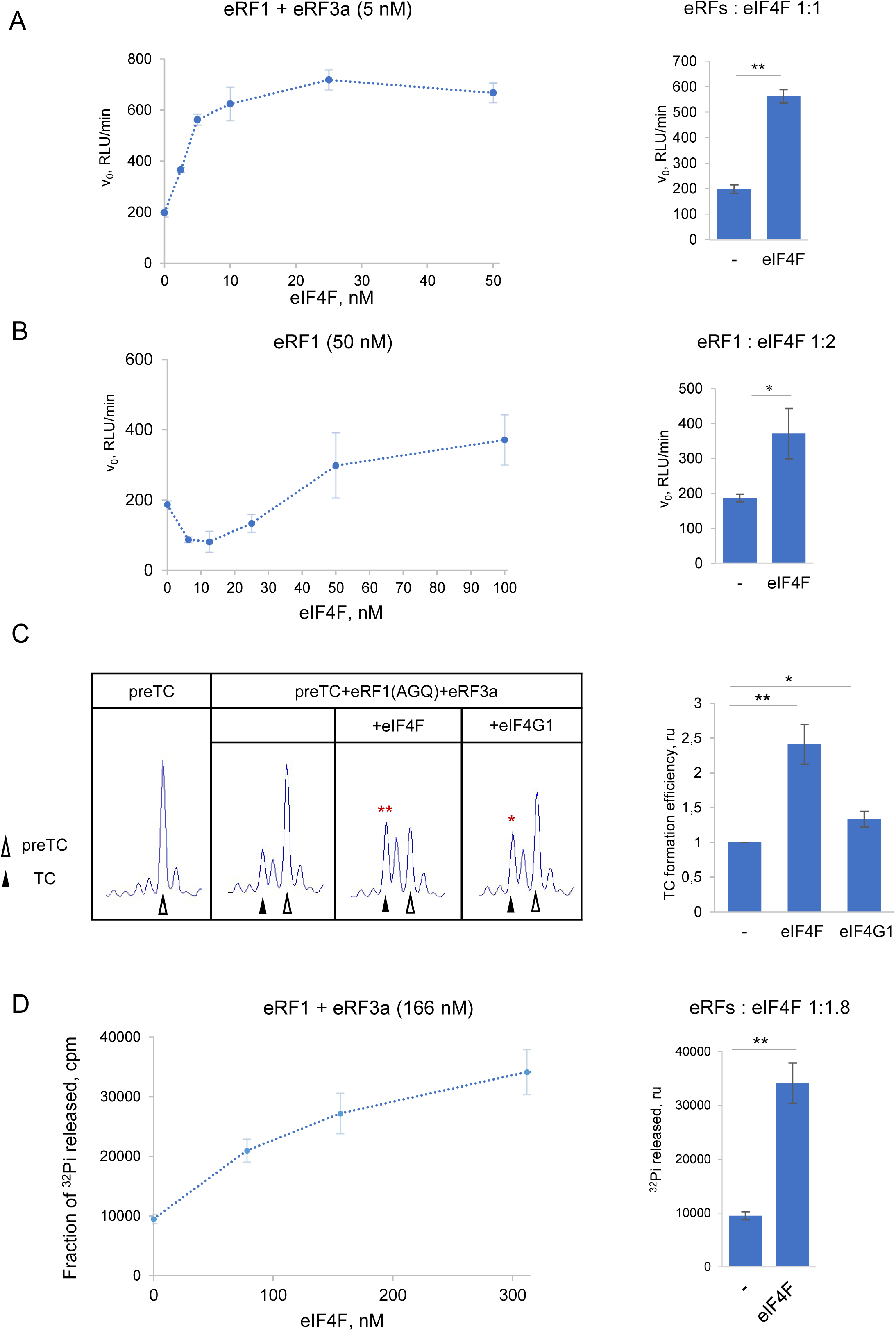
Translation termination is stimulated by eIF4F. (**A**) Rate of peptide release (v_0_) at Nluc preTC induced by eRF1 and eRF3a (5 nM each) in the presence of various concentrations of eIF4F (left) and difference in peptide release rates at selected concentrations: eRFs 5 nM each, eIF4F 5 nM (1:1) (right). (**B**) Rate of peptide release (v_0_) at Nluc preTC induced by eRF1 (50 nM) in the presence of various concentrations of eIF4F (left) and difference in peptide release rates at selected concentrations: eRF1 50 nM and eIF4F 100 nM (1:2) (right). RLU, relative luminescent units. (**C**) Toe-print analysis of TC formation in the reconstituted mammalian translation system in the presence of eRF1(AGQ) and eIF4F or eIF4G1. r.u., relative units. (**D**) GTPase activity of eRF3a in the presence of various concentrations of eIF4F (left) and difference in the amount of the released ^32^P_i_ at selected concentrations: eRFs 166 nM each, eIF4F 312 nM (1:1.8) (right). The data are shown as the mean ± standard error, number of repeats, *n* = 3. Asterisks indicate statistically significant differences between the values (*, *P* < 0.05; **, *P* < 0.01; n.s., not significant).

Translation termination consists of three consecutive stages (see Introduction). To further elucidate the specific stage at which eIF4F exerts its functions, we conducted experiments in a reconstituted *in vitro* translation system ^56^. First, using a fluorescent toe-print assay ^68,69^, we examined the effect of eIF4F on the stop codon recognition by eRF1. Toe-print analysis was performed using preTCs assembled at MVHL-UAA mRNA, which were subsequently purified through equilibrium ultracentrifugation in a sucrose density gradient under high ionic strength conditions. Centrifugation at high ionic strength was necessary to wash preTC from eIFs, since at least eIF3 remains associated with preTC assembled on short CDS in the reconstituted translation system. Increasing KCl concentration during centrifugation to 300 mM facilitated the separation of ribosomes from most eIFs ^61^.

After the accommodation of eRF1, the ribosome protects additional nucleotides on the mRNA, which can be detected in a toe-print assay as a 1-2 nucleotide “shift” of the ribosomal complex ^56^. In this particular experiment, we used the mutant eRF1(AGQ), which binds to the stop codon but is incapable of hydrolyzing peptidyl-tRNA ^70^. Thus, we focused on stages 1 and 2 of translation termination, i.e., the binding of eRF1 to the stop codon and GTP hydrolysis by eRF3, excluding the stage of peptidyl-tRNA hydrolysis. The addition of eIF4F to the reaction significantly promoted preTC to TC transition in the presence of eRF1(AGQ)-eRF3a-GTP (Fig. 1C). Thus, eIF4F promotes either stop codon recognition by eRF1, or GTP hydrolysis by eRF3, or both. Given that eIF4F can stimulate activity of eRF1 even in the absence of eRF3a (Fig. 1B), it appears to stimulate at least the first step of translation termination which can be performed by eRF1 alone.

To address a possibility that eIF4F affects the second step of the termination, we studied if eIF4F alters eRF3a GTPase activity in the presence of eRF1 and ribosomal subunits. Indeed, eIF4F stimulated the GTPase activity of eRF3a by 3.5-fold at 1:1.8 ratio (Fig. 1D). Thus, we can conclude that eIF4F stimulates translation termination both at the stage of eRF1 binding to the ribosome and stop codon recognition, and at the stage of GTP hydrolysis by eRF3a and further accommodation of the GGQ loop in the PTC. However, it cannot be excluded that stimulation of the second stage of termination by eIF4F may be indirect effect resulting from its enhancement of the first stage.

### eIF4G1 and its truncated forms stimulate GTPase activity of eRF3

eIF4F consists of three subunits, i.e., eIF4G1, eIF4A, and eIF4E. The functional activity in translation initiation of reconstructed eIF4F from individual recombinant subunits was confirmed by the assembly of the 48S initiation complex at capped and uncapped mRNAs (Fig. S2). To investigate the contributions of these subunits to translation termination, we assessed the activity of each subunit individually, along with the eIF4G1 homologue eIF4G2 and various truncated forms of eIF4G1 (Fig. 2A). eIF4G1 increased the rate of peptidyl-tRNA hydrolysis by 2.5-fold at a 1:10 ratio of eRFs to eIF4G1 (Fig. 2B, S3A), while in the absence of release factors it did not induce this reaction (Fig. S1C). Thus, individual eIF4G1 also stimulate translation termination, albeit 10 times less efficiently than the complete eIF4F complex. Furthermore, toe-printing also showed that, in the presence of eRF1(AGQ), eIF4G1 stimulated TC formation to a much lesser extent compared to the full eIF4F complex (Fig. 1C). In addition, eIF4G1 failed to stimulate translation termination in the presence of eRF1 alone (Fig. S3B). Therefore, eIF4G1 probably functions at the second stage of translation termination, in which GTP is hydrolyzed by eRF3. Interestingly, eIF4G1 also stimulated activity of the N-terminally truncated eRF3c, lacking the PABP-binding N-domain, and was almost 3-fold more active than full-length eRF3a in the presence of eIF4G1 (Fig. S3C, 2B). Consequently, the activity of eIF4G1 in translation termination was partially suppressed by the N-domain of eRF3. Considering that PABP was present in our system associated with the poly(A) tail, it can be assumed that a complex network of protein-protein interactions governs the interplay among these three proteins in facilitating translation termination.

**Figure 2.**
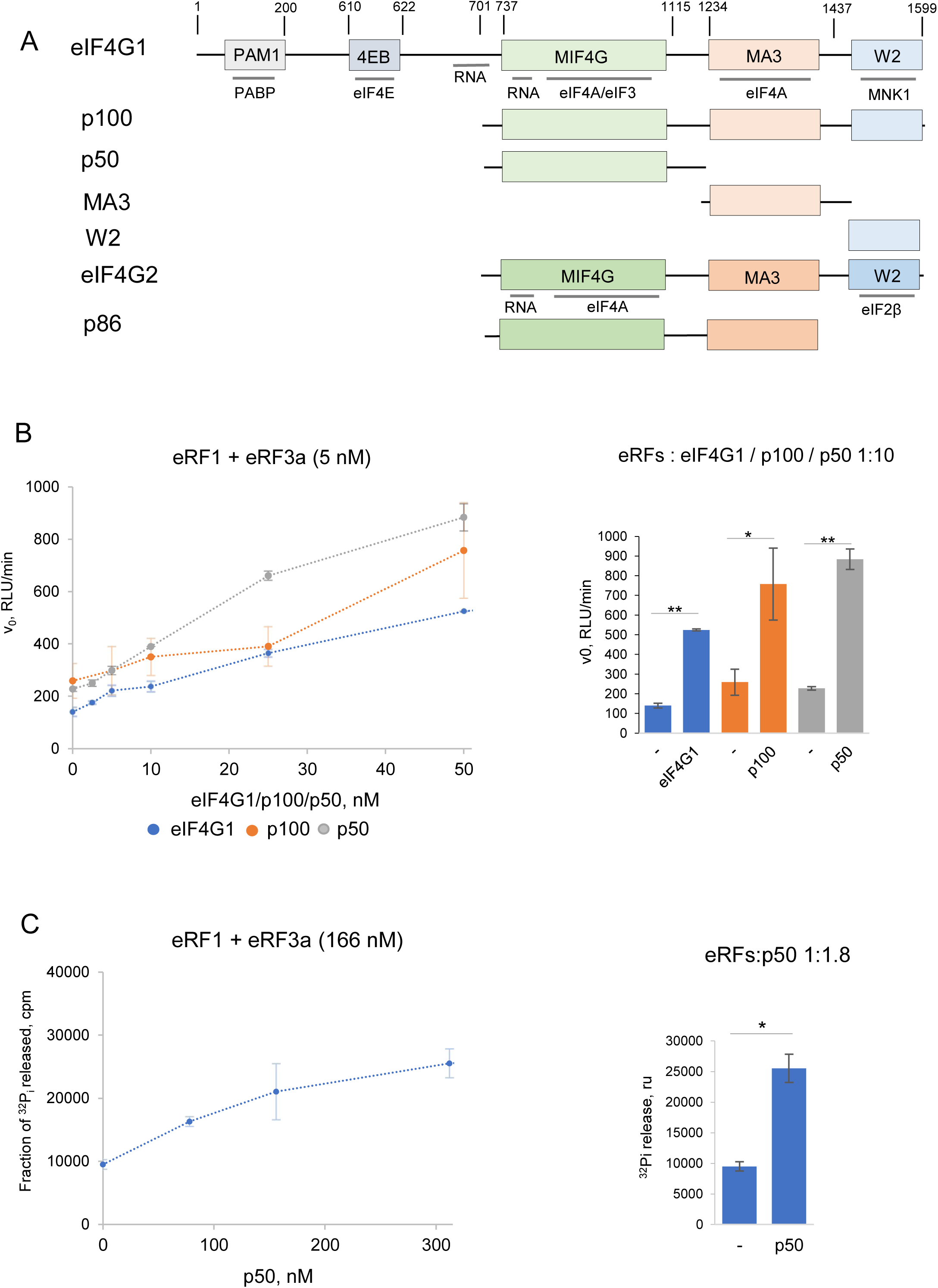
eIF4G and its truncated forms stimulate peptide release and GTPase activity of eRF3. (**A**) Schematic representation of eIF4G, eIF4G2, their truncated forms and domain organization. Regions of interaction with other proteins are indicated. (**B**) Rates of peptide release (v_0_) at Nluc preTC induced by eRF1 and eRF3a (5 nM each) in the presence of various concentrations of eIF4G1, p100, and p50 (left) and difference in peptide release rates at selected concentrations: eRFs 5 nM each, eIF4G1 / p100 / p50 50 nM (1:10) (right). RLU, relative luminescent units. (**C**) GTPase activity of eRF3a in the presence of various concentrations of p50 (left) and difference in the amount of the released ^32^P_i_ at selected concentrations: eRFs 166 nM each, p50 312 nM (1:1.8) (right). The data are shown as the mean ± standard error, number of repeats, *n* = 3. Asterisks indicate statistically significant differences between the values (*, *P* < 0.05; **, *P* < 0.01; n.s., not significant).

Human eIF4G1 consists of five domains that coordinate the translation factors to effectively execute their functions. The spatial arrangement of these domains within eIF4G1, along with the proteins interacting with them, is shown in the scheme (Fig. 2A). To identify the region of eIF4G1 responsible for stimulation of translation termination, we generated truncated forms of the factor: p100, p50, MA3, and W2 (Fig. 2A). The activities of the resulting proteins were tested in the Termi-Luc assay. p100 or p50 stimulated translation termination, and p50 showed even higher activity in translation termination than the full-length eIF4G1 (Fig. 2B). Notably, p100 and p50 lack the PABP-binding region (Fig. 2A). In addition, the p50 variant lacks MA3 and W2 domains, either of which may also be involved in the partial suppression of the activity of full-length eIF4G1 in termination. Importantly, the individual MA3 and W2 domains failed to affect the translation termination (Fig. S4A). In the presence of eRF1 alone, p100 showed some activity in translation termination, which distinguishes it from full-length eIF4G1, which is inactive under these conditions (Fig. S4B).

The activity of p50 was evaluated in a GTPase assay. We found stimulation of eRF3a GTP hydrolase activity by p50, although weaker than by eIF4F (Fig. 2C, 1D). Given that p50 comprises solely the MIF4G domain, we concluded that the GTPase-stimulating activity of eIF4G1 is confined to its MIF4G domain. Therefore, the active forms of eIF4G1 involved in translation termination invariably include the MIF4G domain, and it is the minimum fragment of eIF4G1 required to stimulate translation termination at the second stage - GTP hydrolysis and accommodation of the GGQ loop in the PTC. It is important to note that the N-terminus of eIF4G1, which interactis with PABP in the absence of the eRF3-PABP complex, increases the activity of the release factors in translation termination, and the C-terminus of eIF4G1 partially suppresses the activity of peptide release.

### eIF4G2 acts in translation termination more efficiently than eIF4G1

Like eIF4G1, eIF4G2 contains MIF4G, MA3, and W2 domains, all of which share high similarity with those of the eIF4G1 (Fig. 2A). We obtained recombinant eIF4G2, as well as its truncated form p86 (Fig. 2A), which is generated in cells as a result of proteolytic cleavage during apoptosis. By testing their activity in translation termination using the Termi-Luc assay, we found that both full-length eIF4G2 and p86 stimulated the activity of release factors (Fig. 3A). Interestingly, eIF4G2 was 2.5 times more active than eIF4G1, increasing the rate of hydrolysis by 9-fold at an eRFs:eIF4G2 ratio of 1:10). It is important to highlight that eIF4G2 does not bind PABP (Fig. 2A). While eIF4G2 appears to be an analogue of p100, it exhibits much greater activity in translation termination.

**Figure 3.**
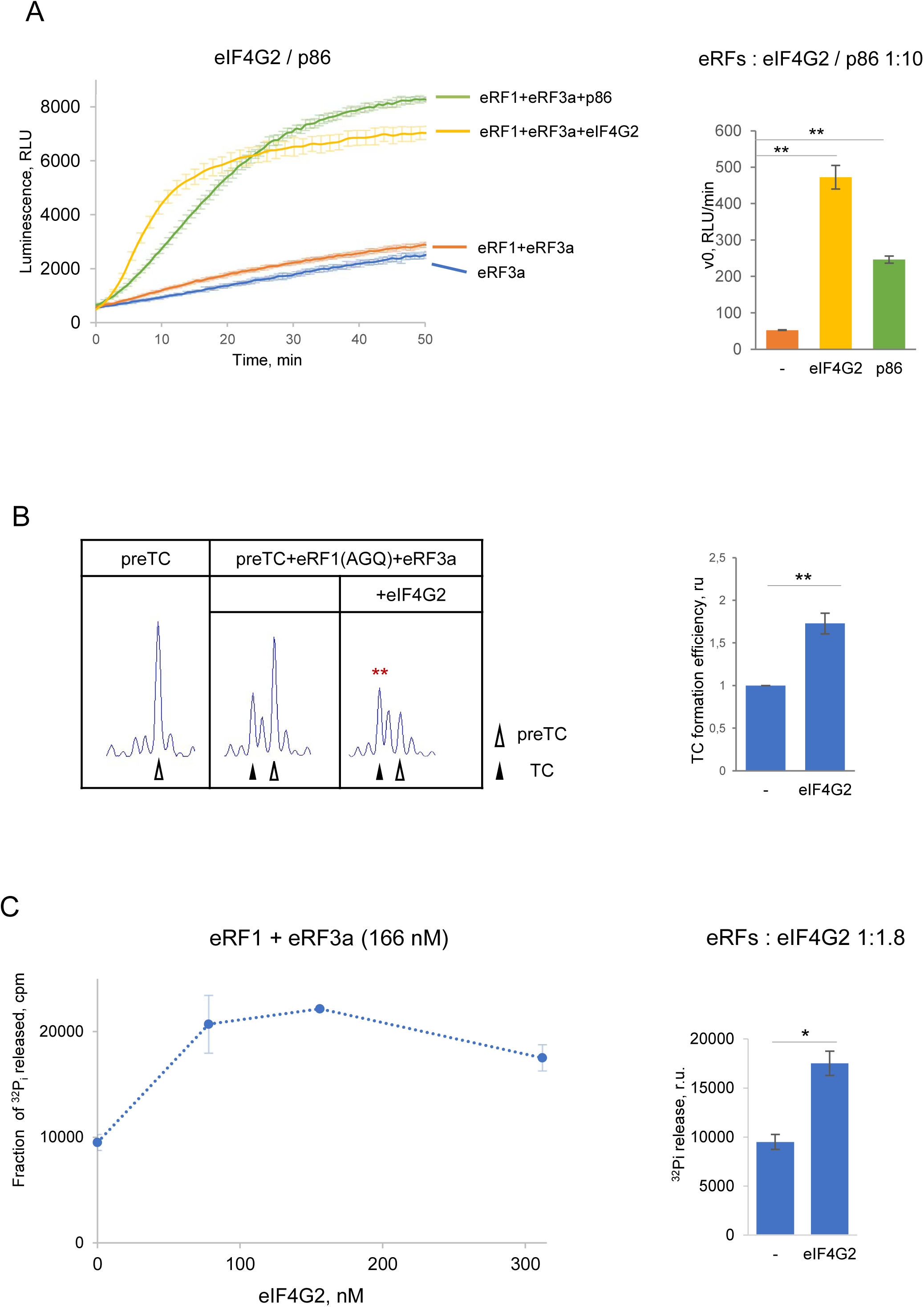
eIF4G2 acts in translation termination more efficiently than eIF4G. (**A**) Nluc release induced by eRF1 and eRF3a in the presence of eIF4G2 and p86 at selected concentrations: eRFs 5 nM each, eIF4G2 and p86 50 nM (1:10) (left) and difference in peptide release rates (right). RLU, relative luminescent units. (**B**) Toe-print analysis of TC formation in the reconstituted mammalian translation system in the presence of eRF1(AGQ) and eIF4G2. r.u., relative units. (**C**) GTPase activity of eRF3a in the presence of various concentrations of eIF4G2 (left) and difference in the amount of the released ^32^P_i_ at selected concentrations: eRFs 166 nM each, eIF4G2 312 nM (1:1.8) (right). The data are shown as the mean ± standard error, number of repeats, *n* = 3. Asterisks indicate statistically significant differences between the values (*, *P* < 0.05; **, *P* < 0.01; n.s., not significant).

The truncated form of eIF4G2, p86, displayed activity comparable to that of p50, increasing the rate of hydrolysis by 4-5 times at an eRFs:p86 ratio of 1:10), which is twice less efficient than the full-length eIF4G2. Given that p86 lacks the W2 domain and that its MA3 domain likely does not affect translation termination (by analogy with that of eIF4G, Fig. S4A), we believe that the W2 domain of eIF4G2 enhances its activity in translation termination. Therefore, W2 domains of eIF4G1 and eIF4G2 differently affect activity of these proteins in translation termination. Subsequent toe-printing and GTPase analyses (Fig. 3B,C) showed that eIF4G2, similarly to eIF4G1, operated at the stage preceding peptidyl-tRNA hydrolysis and stimulated the GTPase activity of eRF3a.

### eIF4A stimulates stop codon recognition by eRF1

The eIF4F complex stimulates translation termination much more efficiently than its core subunit, eIF4G1 (Fig. 1A, 2B). In addition, eIF4G1 only operates in stage 2 of translation termination (Fig. S3B), while eIF4F is active in both stages 1 and 2 (Fig. 1). This suggests that eIF4A and/or eIF4E could also be involved in translation termination. To study their contribution to translation termination, we obtained recombinant eIF4A and eIF4E. The functional activity in translation initiation of purified eIF4A and eIF4E was confirmed by the assembly of the 48S initiation complex in reconstituted translation system at capped mRNA (Fig. S2). The ATP-dependent RNA-helicase activity of eIF4A preparation was confirmed through ATPase assay (Fig. S5). Cap-dependence of the 48S complex assembly in the presence of recombinant eIF4E was confirmed by the assembly at uncapped mRNA (Fig. S2). We then assessed the activities of these proteins in the peptidyl-tRNA hydrolysis reaction in the absence of eIF4G1 using the Termi-Luc assay, using ATP or its non-hydrolysable analogue AMPPNP in the reaction. In the absence of ATP, eIF4A significantly enhanced the release of Nluc from the ribosome in the presence of both release factors, achieving a 5-fold increase in the rate of peptidyl-tRNA hydrolysis at a 1:10 ratio (Fig. 4A, S6A). However, the activity of eIF4A in the presence of both release factors achieving only a 1.5-fold increase in the rate of peptidyl-tRNA hydrolysis at a 1:1 ratio. This is lower than the activity of the entire eIF4F complex in the same conditions (Fig. 1B). Interesingly, eIF4A exhibit a similar stimulating activity in the presence of AMPPNP (Fig. 4A, S6B), while in the presence of ATP the factor stimulated termination only half as efficient (Fig. 4A, S6A). It is possible that during the assay, ATP may have been partially hydrolyzed to ADP, which could subsequently suppress the activity of eIF4A. Therefore, eIF4A promotes translation termination independently of eIF4G1, operating in either an ATP-bound or free form, i.e., in its closed conformation. Conversely, the open conformation resulting from ATP hydrolysis is probably less effective or even inactive in translation termination.

**Figure 4.**
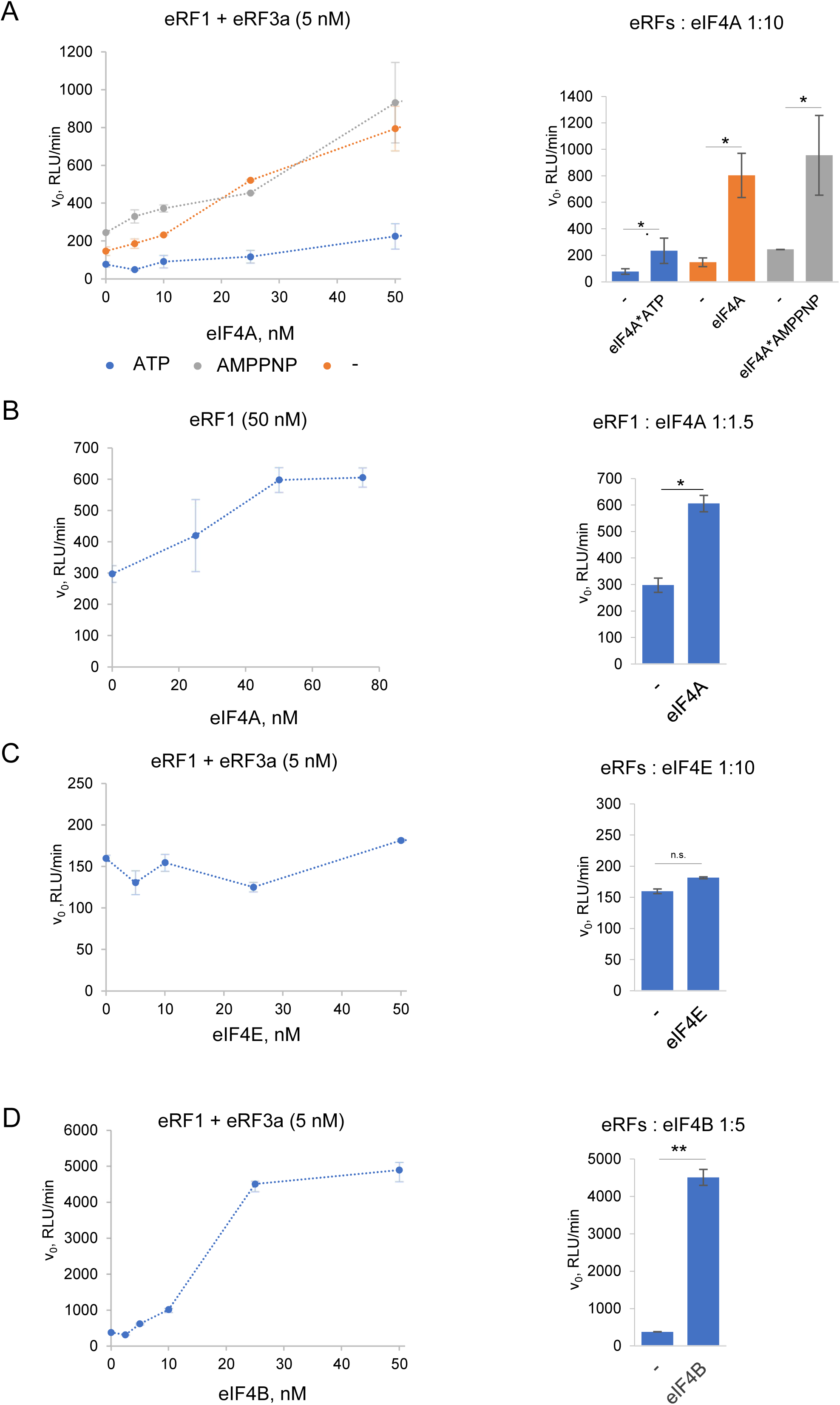
eIF4A, eIF4E and eIF4B activities in translation termination. (**A**) Rates of peptide release (v_0_) at Nluc preTC induced by eRF1 and eRF3a (5 nM each) in the presence of various concentrations of eIF4A (left). eIF4A was pre-incubated with ATP, AMPPNP, or without a nucleotide. The right panel shows difference in peptide release rates at selected concentrations: eRFs 5 nM each, eIF4A 50 nM (1:10). (**B**) Rate of peptide release (v_0_) at Nluc preTC induced by eRF1 (50 nM) in the presence of various concentrations of eIF4A (left) and difference in peptide release rates at selected concentrations: eRF1 50 nM, eIF4A 75 nM (1:1.5) (right). (**C**) Rate of peptide release (v_0_) at Nluc preTC induced by eRF1 and eRF3a (5 nM each) in the presence of various concentrations of eIF4E (left) and difference in peptide release rates at selected concentrations: eRFs 5 nM each, eIF4E 50 nM (1:10) (right). (**D**) Rate of peptide release (v_0_) at Nluc preTC induced by eRF1 and eRF3a (5 nM) in the presence of various concentrations of eIF4B (left) and difference in peptide release rates at selected concentrations: eRFs 5 nM each, eIF4B 25 nM (1:5) (right). RLU, relative luminescent units. The data are shown as the mean ± standard error, number of repeats, *n* = 3. Asterisks indicate statistically significant differences between the values (*, *P* < 0.05; **, *P* < 0.01; n.s., not significant).

In the presence of eRF1 alone, eIF4A, unlike eIF4G1, stimulated peptide release to a degree similar to the complete eIF4F complex, indicating its contribution to the overall eIF4F activity (Fig. 4B, S6C). Since eIF4A activated eRF1 in the absence of eRF3, it is likely that it stimulates the step of translation termination associated with the binding of eRF1 to the stop codon.

We further evaluated eIF4A activity in the presence of the truncated version of the release factor eRF3, eRF3c (Fig. S6D, S7A). We found that eIF4A enhances the efficiency of peptide release to a similar extent as observed in the presence of the full-length eRF3a, suggesting that the activity of eIF4A is independent of eRF3 binding to PABP.

The activity of the remaining subunit of the eIF4F complex, eIF4E, in promoting peptide release was also examined across various concentrations. However, no significant effect was observed (Fig. 4C, S8A). Therefore, eIF4E is unlikely to contribute to the stimulating activity of the eIF4F complex in translation termination.

In summary, our results indicate that two of the three subunits of eIF4F are involved in translation termination, albeit at different steps: eIF4A promotes stop codon recognition by eRF1, while eIF4G1 promotes GTP hydrolysis by eRF3a and the accommodation of the GGQ loop in the PTC.

### eIF4B stimulates translation termination

The activity of eIF4F in translation initiation, particularly that of its subunit eIF4A, is stimulated by eIF4B. It was shown that this factor interacts with both eIF4A ^71–74^, a component of the eIF4F complex, and PABP ^25–28^. It was demonstrated that eIF4B and eIF4G1 cooperate to enhance the ATPase activity of eIF4A (reviewed in ^10,13^). Thus, eIF4B associates with the proteins bound to both the 5’ and 3’ ends of mRNA and may also play a role in translation termination at the closed-loop mRNA structure.

The functional activity of purified eIF4B in translation initiation was confirmed through the assembly of the 48S initiation complex in reconstituted translation system (Fig. S2). Then, we evaluated the activity of purified eIF4B in the peptide release reaction in the presence of eRF1, eRF3a, and PABP (Fig. 4D, S8B). Remarkably, eIF4B exhibited a pronounced stimulatory effect on translation termination, resulting in enhancements exceeding an order of magnitude. Notably, the saturating concentrations of eIF4B required for this effect were lower compared to other individual initiation factors. Specifically, at an eRFs to eIF4B ratio of 1:5, eIF4B increased the rate of peptidyl-tRNA hydrolysis by 12-fold (Fig. 4D). When eRF3a was substituted for eRF3c, the rate of translation termination decreased by approximately two-fold (Fig. S7B, S8C). This suggests that the interaction between eRF3 and PABP may be crucial for the activity of eIF4B in translation termination. Additionally, we found that eIF4B stimulated translation termination in the presence of eRF1 alone, achieving levels comparable to those observed with eIF4A (Fig. S7C, S8D). This indicates that eIF4B may participate in translation termination at the stage of stop codon recognition. However, since its activity significantly increases in the presence of eRF3a, we suppose that it may also be involved in subsequent stages of termination or contribute to overall stability and conformational dynamics of the termination complex.

### eIF4F subunits increase each other’s activity during translation termination

To investigate a mutual influence of eIF4F subunits during translation termination, we performed experiments to assess the stimulation of peptidyl-tRNA hydrolysis by various combinations of these proteins, both in the absence and presence of ATP (Fig. 5). We found that the activity of the combination of eIF4G1 and eIF4A was significantly higher than their individual activities. On the contrary, the addition of eIF4E to the mixture of eIF4G1 and eIF4A resulted in a reduction of the activity (Fig. 5A). This suggests that eIF4E, through its interaction with the capped mRNA, may shift eIF4F activity from termination to the initiation of translation. Notably, as the functional activity of recombinant eIF4E was confirmed in translation initiation (Fig. S2), we assume that inhibitory effect of eIF4E is detected only in translation termination. In the presence of ATP, the coactivation of eIF4G1 and eIF4A was markedly reduced (Fig. 5B), which aligns well with the data obtained earlier for eIF4A in the presence of ATP (Fig. 4A).

**Figure 5.**
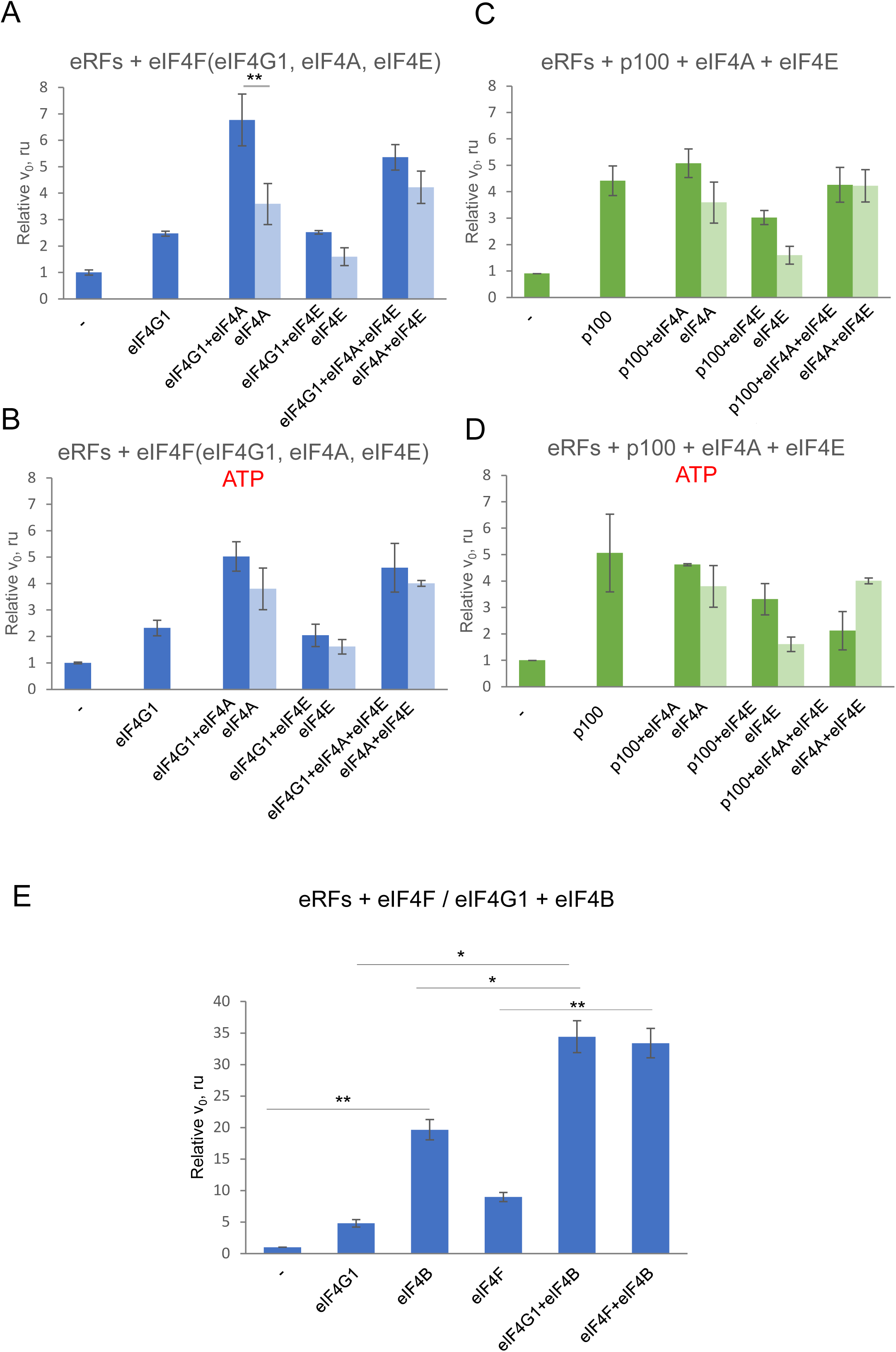
Influence of reconstituted eIF4F on translation termination. Relative peptide-release rates (v_0_) at Nluc preTC induced by eRFs (8 nM each) in the presence of different combinations of eIF4F complex components eIF4G1, eIF4A, eIF4E (35 nM each) (**A**) in the absence of ATP and (**B**) in the presence of ATP. p100, eIF4A, and eIF4E (20 nM each) (**C**) in the absence of ATP and (**D**) in the presence of ATP. (**E**) Relative peptide-release rates (v_0_) at Nluc preTC induced by eRFs (8 nM each) in the absence and presence of eIF4B (15 nM) and eIF4G1 (35 nM) / eIF4F (5 nM). The data are shown as the mean ± standard error, number of repeats, *n* = 3. Asterisks indicate statistically significant differences between the values (*, *P* < 0.05; **, *P* < 0.01; n.s., not significant).

To further elucidate the influence of distinct regions of eIF4G1 on the activity of reconstituted eIF4F in translation termination, we compared the results obtained for eIF4G1 with those for p100. This comparison revealed no synergistic interaction between p100 and eIF4A in translation termination (Fig. 5C,D). Although p100 contains eIF4A-binding domains, its conformation appears to preclude eIF4A from exerting a synergistic effect within the complex. Furthermore, the addition of eIF4E, which lacks a binding site for p100, slightly diminished its stimulatory activity both in the presence and absence of eIF4A (Fig. 5C), suggesting some slight non-specific inhibitory effect of eIF4E on translation termination in this context.

Additionally, we investigated the potential mutual influence of the factors eIF4F/eIF4G1 and eIF4B during translation termination. Combinations of these factors exhibited greater activity than the individual proteins, indicating probably additive effects (Fig. 5E). This suggests that the mechanisms by which eIF4F and eIF4B stimulate translation termination operate independently. Interestingly, eIF4B equally well stimulated both the eIF4F ternary complex and eIF4G1 alone.

In conclusion, our findings indicate that the eIF4G1-eIF4A-eIF4B complex, or their combination, is more active in translation termination than its individual components, with the N-terminal region of eIF4G1 being essential for it to function full-throttle. This region is capable of binding both PABP and eIF4E, but inasmuch as eIF4E does not promote the activity of the eIF4G1-eIF4A complex, we tend to think that it is PABP, associated with the poly(A) tail of the mRNA, that plays a crucial role in the eIF4F-mediated enhancement of translation termination. Furthermore, as eIF4B can also bind to PABP, it is possible that the co-stimulation by eIF4G1-eIF4B may be mediated by the poly(A) tail as well.

### Poly(A) tail of mRNA increases stimulation of translation termination by eIF4G1, eIF4A, and eIF4B

To assess the effect of PABP bound *in cis* to the poly(A) tail on the activities of eIF4G1, eIF4A, and eIF4B during translation termination, we assembled preTCs in RRL on both polyadenylated and non-polyadenylated mRNAs, as well as capped and uncapped mRNAs, and purified them through sucrose gradient centrifugation. At a ratio of 2:5 to eRFs, eIF4F effectively stimulated peptide release across all tested mRNAs (Fig. S9A-D), exhibiting slight variations among the four conditions, ranging from 1.5 to 3-fold (Fig. S9E). Therefore, eIF4F is capable of activating translation termination in both the presence and absence of PABP.

We further assayed the potential dependence of the activity of eIF4B and the individual subunits of eIF4F on the presence of a poly(A) tail in translation termination (Fig. 6). Each of the three factors – eIF4G1, eIF4A, and eIF4B – exhibited enhanced stimulation of translation termination in the presence of the poly(A) tail, while eIF4E, as expected, showed no effect on peptide release regardless of the presence of the poly(A) (Fig. 6). The effect of the poly(A) tail on the activity of eIF4B and eIF4G1 can be attributed to their direct interactions with PABP. In contrast, eIF4A, which does not directly interact with PABP, may exert its poly(A) tail-dependent stimulatory activity through a complex network of interactions involving PABP, eRF3a, and eRF1. As we have shown above, eIF4A promotes the ribosome loading of eRF1, which also depends on eRF3a and PABP.

**Figure 6.**
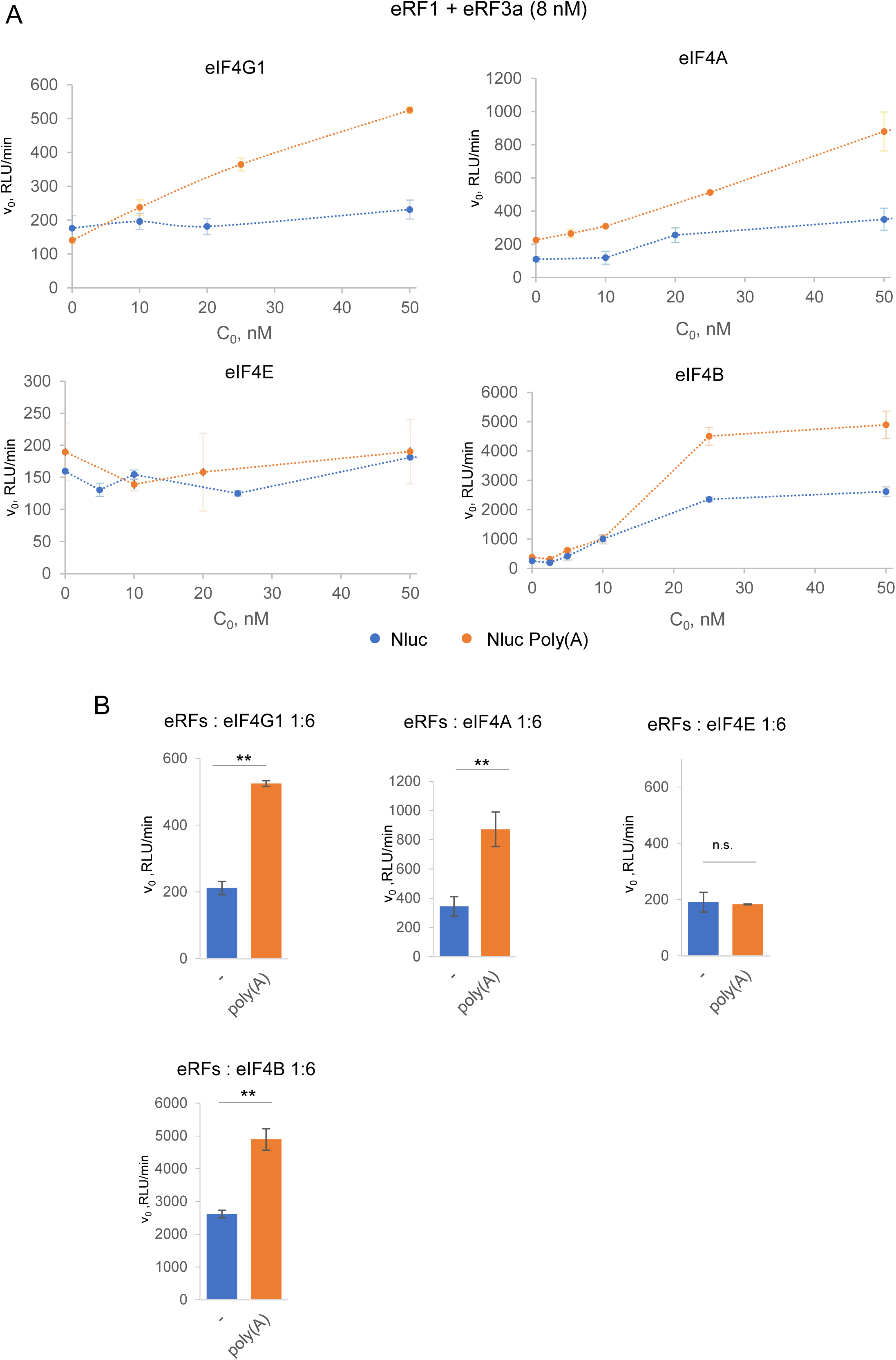
Effect of poly(A) tail, bound with PABP, on the activity of eIF4F components in translation termination. (**A**) The rates of peptide release (v_0_) at preTCs assembled on Nluc mRNA without poly(A) tail (the blue curves) or with poly(A) tail (the orange curves). Peptide release was induced by eRF1 and eRF3a (8 nM each) in the presence of various concentrations of eIF4G1, eIF4A, eIF4E, and eIF4B. (**B**) Differences in peptide release rates at selected concentrations: eRFs 8 nM each, eIF4G1, eIF4A, eIF4E, and eIF4B 50 nM (1:6). RLU, relative luminescent units. The data are shown as the mean ± standard error, number of repeats, *n* = 3. Asterisks indicate statistically significant differences between the values (*, *P* < 0.05; **, *P* < 0.01; n.s., not significant).

The results presented here clearly demonstrate that while PABP enhances the function of eIF4F and eIF4B in promoting translation termination, it is not an obligatory factor in this process.

### eIF4A promotes the loading of eRF1 on the ribosome, while eIF4G1 enhances the dissociation of eRF1-eRF3a from the ribosome

The involvement of the eIF4F complex in translation termination implies that its subunits may physically interact with release factors or the terminating ribosome, thereby stimulating their activities. To elucidate how initiation factors participate in translation termination, we examined if these proteins bind preTC or the release factors.

Since the interaction of the eIF4F complex with the terminating ribosomes has not been studied previously in a reconstituted system, we conducted a series of experiments to assess the binding of eIF4A, eIF4G1, and p100 to preTCs. For this purpose, preTCs were assembled on the MVHL-UAA mRNA in the reconstituted mammalian translation system and subsequently purified using equilibrium ultracentrifugation in a sucrose density gradient under high ionic strength conditions. The purified preTCs were then incubated with eIF4F subunits, followed by a second round of centrifugation at 100 mM KCl. Proteins from the gradient fractions were precipitated and analyzed via Western blotting using specific antibodies.

In the presence of GTP, the release factors were barely detectable in the ribosomal fractions. This observation suggests that following GTP hydrolysis, they likely dissociate from the ribosomes during the second round of ultracentrifugation, as no cross-linking agents were used to stabilize protein interactions in these experiments (Fig. 7A,D). However, eIF4A was found in preTC when AMPPNP was included, but was nearly undetectable when ATP was present (Fig. 7B,D). This suggests that eIF4A interacts with preTC in the ATP/AMPPNP-bound form, while subsequent ATP hydrolysis promotes its dissociation from the ribosomes. When both eRFs and eIF4A were added to preTC, we observed an increase in the amount of eRF1 associated with the ribosomal complex in the presence of both ATP and AMPPNP (Fig. 7C,D). Importantly, in the presence of eRFs, eIF4A dissociated from the ribosome even in the presence of AMPPNP (Fig. 7C, D), which confirms the authentic binding of eIF4A to preTC. Notably, eRF3a was not detected in the ribosomal complex even in the presence of eIF4A. We suppose that eIF4A binding to a preTC in an ATP-bound state promotes the loading of eRF1 into the ribosome, after which eIF4A hydrolyzes ATP and dissociates from the ribosome in an ADP-bound form. Control experiments without ribosomes ruled out potential oligomerization of the proteins under the reaction and centrifugation conditions (Fig. S10A).

**Figure 7.**
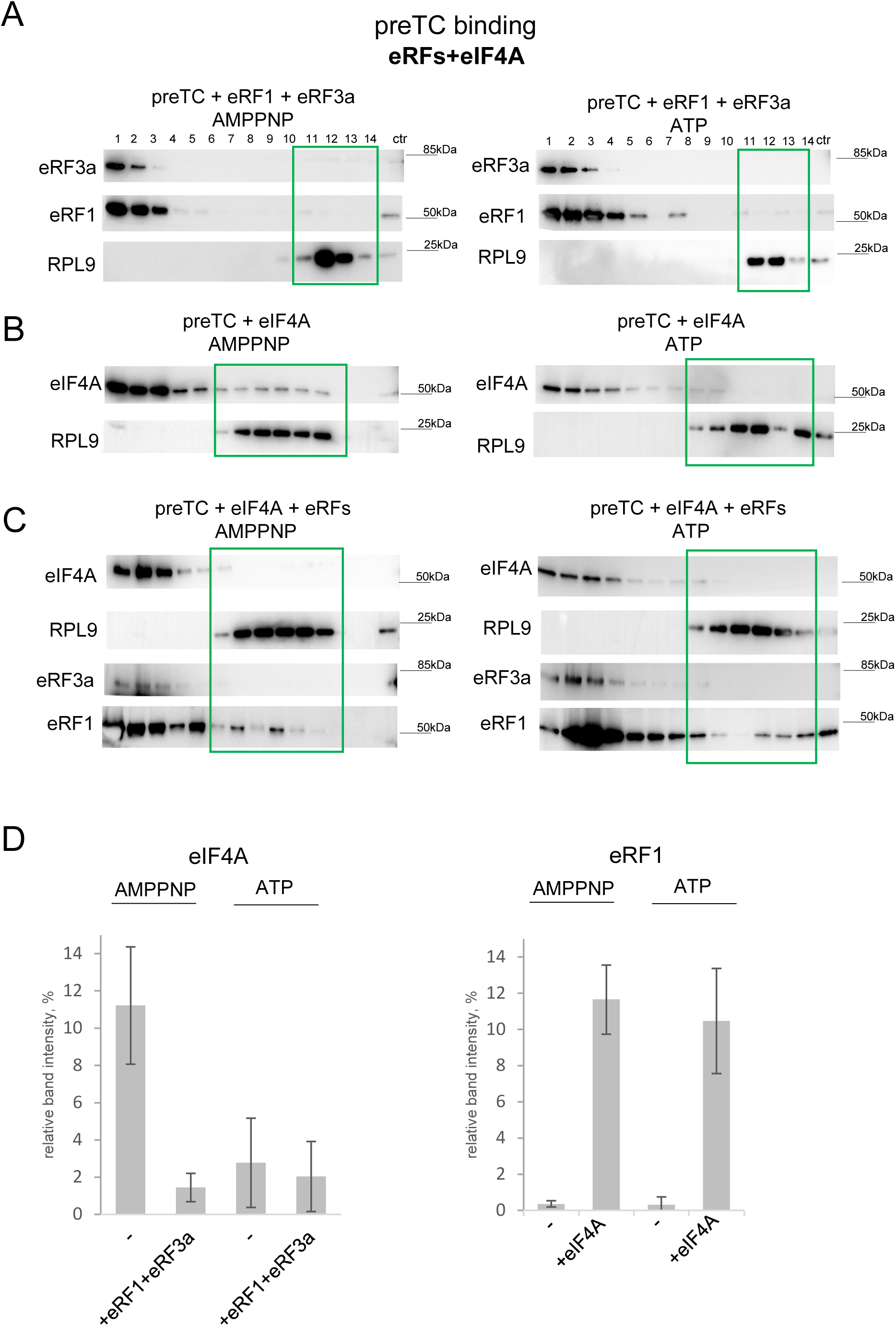
eIF4A promotes the loading of eRF1 to the ribosome. (**A**) Western blot analysis of SDG fractions obtained after incubation of preTC with eRF1 and eRF3a in the presence of ATP or AMPPNP. (**B**) Western blot analysis of SDG fractions obtained after incubation of preTC with eIF4A in the presence of ATP or AMPPNP. (**C**) Western blot analysis of SDG fractions obtained after incubation of preTC with eRFs and eIF4A in the presence of ATP or AMPPNP. Fraction numbering: from 1 to 14, from top to bottom of SDG. PreTC-associated fractions are marked with a green frame. Antibodies raised against the proteins of interest were used. (**D**) Relative band intensities of eIF4A (the left panel) or eRF1 (the right panel) in the preTC-associated fractions. Number of repeats: n = 2 (for AMPNP), n = 3 (for ATP).

Since individual eIF4A and eIF4G1 both are able to independently operate in translation termination, we also performed experiments on the binding of eIF4G1 or p100 to the purified preTCs or TCs. We found that eIF4G1 is unstable under these conditions: it likely aggregated during the incubation and then precipitated, as confirmed by control ultracentrifugation of eIF4G1 without ribosomal complexes. p100, on the other hand, did not bind stably to preTC. However, we were able to detect small quantities of release factors associated with the ribosomes, which allowed us to ascertain that p100 promotes the dissociation of the eRF1-eRF3a complex from the ribosome in the presence of GTP, while not affecting the dissociation of individual eRF1 (Fig. S10B). In the presence of a non-hydrolysable analogue of GTP, GDPCP, no binding of the release factors to the ribosome could be detected. Thus, we have shown that p100 promotes the dissociation of release factors from the ribosome after GTP hydrolysis, acting specifically in the presence of eRF3a. This is consistent with our previous observations of the stimulatory activity of eIF4G1 in the second stage of translation termination during GTP hydrolysis by eRF3a (see Fig. 2).

### Release factors are unable to bind eIF4F or eIF4B in solution

Next, we tested binding of the release factors to His-tagged eIF4A, eIF4E, eIF4B, and eIF4G1 domains (p100, p50, and MA3). The release factors without the His-tag were mixed with the proteins of interest, which were pre-bound to an affinity resin. After incubation and washes, bound complexes were eluted with imidazole and analyzed by Western blotting with antibodies against release factors or His-tag. The acetate kinase (Ack) from *E. coli*, which also carried His-tag, was used as a negative control. Neither eIF4A nor eIF4B interacted with eRF1 and eRF3a in solution, since the amounts of factors eluted from the putative complexes were comparable to those in the negative control (Fig. S11A). Thus, we could only determine the nonspecific binding of eRFs to the resin.

To evaluate the binding interactions between eIF4E, eIF4G1, and its domains with release factors eRF1 and eRF3a, we conducted a series of co-precipitation assays using recombinant His-SUMO-tagged release factors and His-tagged eIF4G1. Our results demonstrated that neither eIF4E, nor full-length eIF4G1 or its individual domains exhibited binding to the release factors in solution (Fig. S11B,C and S12). Importantly, we found no detectable interactions between eIF4G1 domains and eRF3a, even in the presence of GTP, GDP, or GDPCP in the reaction mixtures. These findings suggest that the initiation factors investigated do not interact directly with the release factors in solution. Rather, it appears that their role in translation termination is mediated through modulation of the release factors’ activity on the ribosome.

## DISCUSSION

Upon the formation of the mRNA closed-loop structure, the cap-binding initiation complex located at the mRNA 5’ end is brought into the close proximity with the termination complex due to the simultaneous interaction of PABP, which is bound to the poly(A) tail of mRNA, with initiation factor eIF4F and release factors. In this study, we explored the role of eukaryotic translation initiation factors, particularly eIF4F, in the process of translation termination in mammals. The results regarding the activities of different initiation factors and their truncated forms at various stages of translation termination are summarized in Table 1.

**Table 1.**
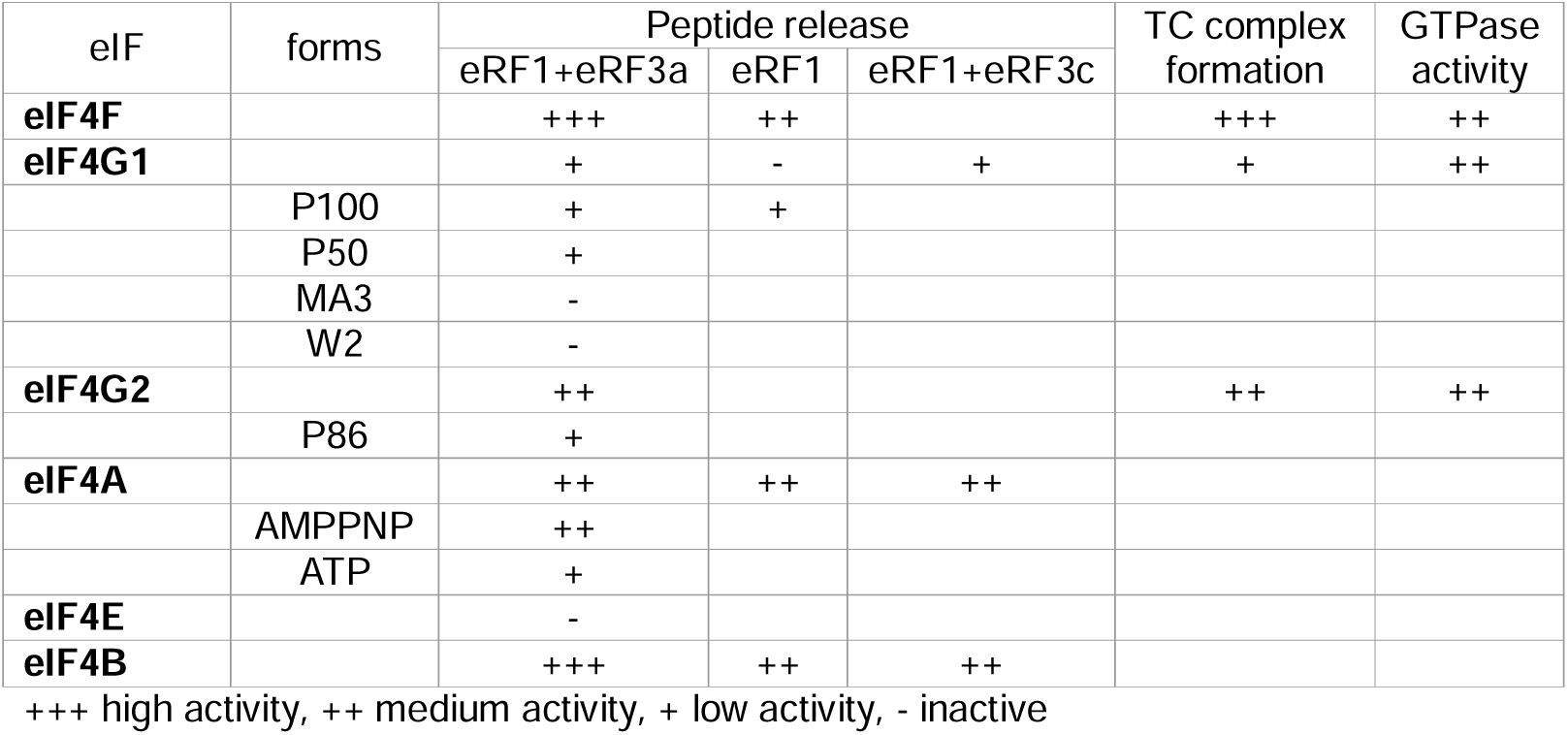
Activities of eIFs in different stages of translation termination.

| eIF | forms | Peptide release |  |  | TC complex formation | GTPase activity |
| --- | --- | --- | --- | --- | --- | --- |
|  |  | eRF1+eRF3a | eRF1 | eRF1+eRF3c |  |  |
| **eIF4F** |  | +++ | ++ |  | +++ | ++ |
| **eIF4G1** |  | + | - | + | + | ++ |
|  | P100 | + | + |  |  |  |
|  | P50 | + |  |  |  |  |
|  | MA3 | - |  |  |  |  |
|  | W2 | - |  |  |  |  |
| **eIF4G2** |  | ++ |  |  | ++ | ++ |
|  | P86 | + |  |  |  |  |
| **eIF4A** |  | ++ | ++ | ++ |  |  |
|  | AMPPNP | ++ |  |  |  |  |
|  | ATP | + |  |  |  |  |
| **eIF4E** |  | - |  |  |  |  |
| **eIF4B** |  | +++ | ++ | ++ |  |  |

+++ high activity, ++ medium activity, + low activity, - inactive

Our findings indicate that eIF4F effectively promotes translation termination. We found that individual subunits of the eIF4F complex stimulate the activity of different release factors: eIF4A promotes loading of eRF1 into the ribosome, while eIF4G1 enhances the GTPase activity of eRF3a. Notably, our data demonstrate that both eIF4G1 and eIF4G2 exhibit this activity in translation termination, even despite eIF4G2 does not bind PABP. Importantly, we identified MIF4G domain as the minimal functional unit capable of stimulating translation termination. Furthermore, our findings extend to eIF4B, another initiation factor associated with PABP, which also displays a stimulatory effect on translation termination.

eIF4A has recently been shown to bind to the 48S complex at two sites, one of which is located near the mRNA entry channel ^73^. This may also be the site of eIF4A binding to the 80S ribosome, as our findings indicate that eIF4A can interact with the preTC and, modulate eRF1 binding to the A site (Fig. 7). Only a minor portion of the eIF4G1 is resolved in the latest cryo-EM structures of the human 48S complex, raising questions about whether the second eIF4A molecule is recruited to the mRNA entry by the MA3-domain of eIF4G1 or if it binds independently of eIF4G1.

It has previously been shown that the helicase involved in the transport of mRNA from the nucleus, DDX19 in human (Dbp5 in yeast), is also involved in translation termination ^59,60^, facilitating the loading of eRF1 into the ribosome ^60^. Notably, DDX19/Dbp5 shares structural similarities with eIF4A. In its ATP-bound form, DDX19 interacts with preTC and, likely by changing the conformation of the A site, enhances the binding of eRF1 to the stop codon. Following ATP hydrolysis, DDX19 subsequently dissociates from the ribosome ^60^. In the currents study, we show a similar stimulatory effect on termination by eIF4A (Fig. 4, 7), which implies that the two proteins may perform analogous functions in this process. While the precise role of DDX19 in the general mechanism of translation termination remains unclear, it is plausible that, together with Gle1, its partner in mRNA transport from the nuclear pore and structural homologue of eIF4G1, these proteins could functionally substitute for eIF4F in translation termination.

Other proteins that share structural homology with eIF4G1, which have a hand both in translation initiation and termination, include PDCD4 and PAIP1. Tumor suppressor PDCD4, a multifunctional protein, inhibiting cell growth, tumor invasion, metastasis, and inducing apoptosis ^75^ contains two MA3 ^76^ domains, which bind to eIF4A, and the N-terminal domain of this protein interacts with PABP ^77^. We showed previously that PDCD4 binds to preTC and stimulates translation termination via promoting the GTPase activity of eRF3 and its subsequent dissociation from the ribosome ^64^. Therefore, PDCD4 may serve as a functional analogue of eIF4G1 in the context of translation termination, with the potential to displace it from the termination complex through interactions with preTC and PABP. PAIP1 contains a MIF4G domain homologous to that of eIF4G1 ^66^, and this protein, as its name implies, also binds to PABP. PAIP1 facilitates the formation of termination complexes in the presence of eRF3 and binds to the eRF3-eRF1 complex but does not stimulate the hydrolysis of peptidyl-tRNA and inhibits the activity of PABP in translation termination ^66^. We proposed that this protein can reduce the efficiency of translation termination via binding with PABP in solution. Therefore, while the MIF4G domain of PAIP1 retains some functional properties of its eIF4G1 counterpart, it lacks the capacity to stimulate the GTPase activity of eRF3. This feature allows PAIP1 to function as an inhibitor of translation termination in some instances, e.g. at premature termination codons, or under some conditions. Collectively, these data suggest that the mechanism we have described for the stimulation of translation termination by eIF4F is fundamentally conserved and is frequently used by the cell as a target for regulating various processes, including readthrough of premature termination codons, translation pausing, and the induction of apoptosis under stress.

The eukaryotic initiation factor eIF4B interacts with eIF4A, as well as with the ribosome, to promote translation initiation ^10,13,71–74,78^. However, in yeast, deletion of eIF4B reduces the relative translation efficiencies of many more mRNAs than does inactivation of eIF4A ^17^. This might be related to the activity of eIF4B in translation termination, as we have shown in this study (Fig. 4D). According to recent structural data ^73^, eIF4B is bound in the vicinity of the mRNA entry channel and thus unlikely can interact with release factors directly. However, it may contribute to translation termination through enhancing the eIF4A activity and by stabilizing the termination complex indirectly by changing the conformation of the 40S subunit.

The minimal domain of eIF4G1 that is active in translation termination, MIF4G (Fig. 2B), is capable of interacting with eIF4A, eIF3, and RNA. eIF4G2, which also promotes the termination (Fig. 3), binds eIF4A and, likely, eIF3, but not eIF4E or PABP. In addition, eIF4G1 and eIF4A stimulate translation termination in the presence of eRF3c, which is unable to bind to PABP (Fig. S3C, S7A). Therefore, presence of PABP and eIF4E is not required for the eIF4F complex to promote translation termination. However, we also showed that, unlike the full-length eIF4G1, p100 does not increase the activity of eIF4A in termination (Fig. 5A, C). This means that the N-terminal region of eIF4G1 is important for efficient activation of translation termination. In addition, the presence of a poly(A) tail increases the activities of all tested eIFs in termination (Fig. 6). These observations point to a more intricate regulatory mechanism governing translation termination, which encompasses the individual contributions of eIF4A and eIF4G1, as well as their interactions with one another. Importantly, these interactions are likely mediated by the poly(A) tail-bound PABP and the ribosome, suggesting a multifaceted framework for the regulation of translation termination.

Based on the obtained data, we propose a comprehensive model for the eIF4F and eIF4B function in translation termination (Fig. 8). In this model, PABP, which is bound to the poly(A) tail of the mRNA and may be represented by multiple molecules, serves as the platform for the assembly of several factors involved in translation termination. eIF4F and eIF4B associate with PABP molecule(s), resulting in the formation of a closed-loop mRNA structure. The eRF1-eRF3-GTP complex then associates with either the same or a neighboring PABP molecule on the poly(A) tail. As the elongating ribosome encounters the stop codon and stalls while waiting for eRF1, eIF4A and eIF4B from the nearby PABP-eIF4F complex bind to the 40S subunit and induce its conformational changes. PABP promotes the binding of eRF3-GTP to the pretermination complex, while ATP-bound eIF4A accelerates the loading of eRF1 into the A site. eIF4B stimulates these processes and stabilizes the optimal ribosome conformation. The MIF4G domain of eIF4G1 enhances the GTPase activity of eRF3a on the ribosome. As a result, the release factor complex undergoes conformational changes, accommodating the GGQ loop of eRF1 in the PTC. This induces the hydrolysis of peptidyl-tRNA and the release of the synthesized peptide from the ribosome. Subsequently, eIF4G1 promotes the dissociation of release factors from the ribosome, while eIF4A hydrolyzes ATP and also dissociates. Notably, eIF4A is the most abundant eukaryotic translation initiation factor, suggesting that it may participate in processes beyond initiation. It is possible that eIF4F subunits reassemble into a complex on PABP, enabling their further participation in both translation termination and initiation. After translation termination, eRFs may also bind to PABP awaiting the next ribosome to arrive at the stop codon. It is also important to note here that cap-bound eIF4E in the initiation complex appears to switch the activity of the other two eIF4F subunits between participation in initiation and termination.

**Figure 8.**
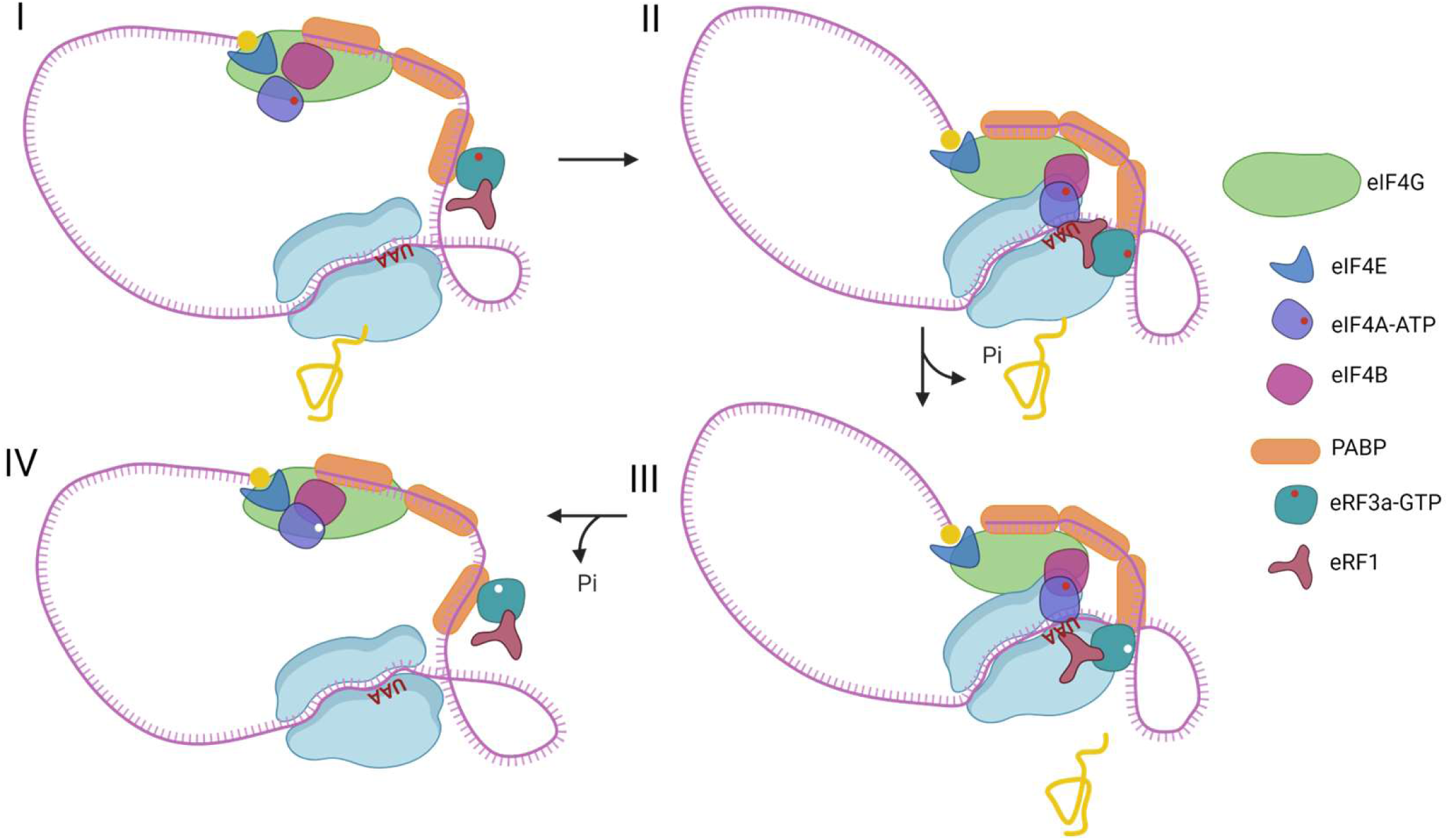
The model of eIF4F functioning in translation termination. The figure is created with BioRender.com.

In addition to the eIF4F molecules recruited to the termination complex by the poly(A)-bound PABP, translation termination may also be affected by eIF4F present in the solution. However, PABP facilitates the interaction of all the participants at once by bringing initiation factors closer to the terminating ribosome, thereby increasing the likelihood of rapid translation termination (Fig. 6). In addition, data regarding the activity of truncated forms of eIF4G1 and eRF3 (Fig. 2, S3) indicate that these proteins compete for interaction with PABP with their N domains, although they bind to different regions of PABP (Fig. 2A). Thus, the activity of full-length eIF4G1 is diminished compared to that of p100, as well as the activity of eRF3a is reduced compared to that of eRF3c. This competition was not observed in the presence of eIF4A as a part of the eIF4F complex, likely due to a conformation change in eIF4G1. Similarly, the high activity of eIF4G2 in translation termination may result from a lack of competition with eRF3a for binding to PABP, as eIF4G2 does not have a PABP binding site (Fig. 3).

eIF4G1 has long been implicated in reinitiation on mRNA with uORFs ^79,80^. Both eIF4G1 and eIF4A present in elongating and terminating ribosomes on reinitiation-permissive uORFs in yeast ^81^, while eIF4G1 is retained on elongating ribosomes in human ^82^. Considering that eIF4G2 can substitute for eIF4G1 and that it is involved in scanning and reinitiation after translation of uORFs ^36,37^, we hypothesize that eIF4G2 may also directly participate in translation termination at the uORFs’ stops. eIF4G2 interacts with eIF4A ^83^ and thus may form an eIF4F-like complex capable of affecting translation termination. In line with this notion, the mutant eRF1(AGQ) suppressed eIF4G2-dependent translation of both uORF and subsequent CDS ^37^. The binding of eIF4G2 to the terminating ribosome is probably enhanced by the W2 domain, since the truncated form of this protein demonstrates a twofold reduction in stimulation of translation termination compared to the full-length protein (Fig. 3A). Collectively, our data suggest eIF4G1 and eIF4G2 not only facilitate the resumption of ribosome scanning, but also promote efficient termination, which may be a prerequisite for efficient reinitiation.

Apart from viral infection, eIF4G1 and eIF4G3 are also cleaved by caspase-3 to a 76-kDa fragment (M-FAG) that lacks PABP-binding site and MA3-W2 domains (reviewed in ^84^), and eIF4G2 is also cleaved by caspase-3 to remove its W2 domain ^85^. The differential activities of eIF4G1 fragments described here suggest that apoptotic and viral reprogramming of the translational machinery may not be limited to translation initiation, but may affect translation termination as well.

We would like to emphasize that the termination complex associated with the closed-loop mRNA structure, featuring eIF4F and eIF4B, may encompass additional factors. For example, we have previously demonstrated that eIF3j facilitates the loading of release factors into the ribosome ^61^. Additionally, the yeast ribosome recycling factor Rli1 has been shown to positively affect the rate of peptidyl-tRNA hydrolysis ^58^. Understanding the intricate interactions among these proteins during the termination process may serve as an important avenue for future investigations.

Finally, effective translation termination is also a critical issue for mRNA stability, particularly in preventing nonsense-mediated decay (NMD) ^86^. The role of PABP in inhibiting NMD is well-established (reviewed in ^87,88^). Notably, some studies have also highlighted the significant involvement of eIF4G in regulating this process. A series of investigations conducted by the Romão group demonstrated that mRNAs harboring premature termination codons in close proximity to the translation initiation codon can evade NMD through a mechanism that involves the interaction of PABP with the translation initiation complex (see ^89^ and references therein). This finding aligns with the established notion that NMD majorly tolerates translation termination at the stop codon of uORFs, despite the fact that such stop codons may be recognized as premature termination codons according to NMD principles. Furthermore, in 2014, two independent research groups reported that the interaction between PABP and eIF4G is essential for suppressing NMD, and that the direct tethering of eIF4G to an NMD reporter is sufficient to antagonize NMD factor activity ^90,91^. The authors proposed that eIF4G may play a role in both translation termination and NMD suppression. Additionally, eIF4B was recently shown to interact with UPF1, the key NMD factor ^92^. We suggest that it is the close proximity to initiation complex that allows both authentic CDS stop codons and uORF ones to evade NMD.

In summary, our results suggest that during the formation of the closed-loop mRNA structure, the translation rate increases not only due to stimulation of initiation but also due to stimulation of termination, with the factors that form the closed-loop structure, eIF4F and PABP, playing critical roles in both processes.

## ACKNOWLEDGEMENTS

We are grateful to Ludmila Frolova for providing us with plasmids encoding eRF1, eRF1(AGQ) and eRF3c, to Tatyana Pestova and Christopher Hellen who provided us with plasmids encoding translation initiation factors, to Boris Eliseev and Christian Schaffitzel who provided us with plasmid encoding eRF3a and for assistance with the expression of eIF4F, eIF4G and eRF3a in Baculovirus expression system. Sequencing of plasmids and cDNA fragment analyses were performed by the center of the collective use “Genome” of EIMB RAS. We are thankful to the Centre for Precision Genome Editing and Genetic Technologies for Biomedicine for access to the facilities necessary for this study. The graphical abstract is created with BioRender.com. The study was supported by the Russian Science Foundation (Grant No. 22-14-00279).

## AUTHOR CONTRIBUTIONS

E.S., A.S., W.S., A.I., N.B. and T.E. conducted the experiments; E.A. designed the experiments, E.A, S.D. and I.T. wrote the paper.

## SUPPLEMENTAL INFORMATION

Supplementary figures. Figures S1–S12. Supplementary data. Supplemenary figure legends.

## Methods

### Ribosomal subunits and translation factors

The 40S and 60S ribosomal subunits and eukaryotic translation factors eIF2, eIF3, eEF1H, and eEF2 were purified from RRL and HeLa cell lysates as described previously ^56^. The human translation factors eIF1, eIF1A, eIF4A, eIF4B, p50 (MIF4G domain of eIF4G1), ΔeIF5B, eIF5, eRF1, eRF1(AGQ) variant, and eRF3c lacking the N-terminal domain (138 amino acid residues including PAM2 domain) were expressed in *E. coli* strain BL21(DE3) and purified via Ni-NTA agarose followed by ion-exchange chromatography ^56^. Human eRF3a was expressed in insect cells Sf21 using the EMBacY baculovirus from the MultiBac Expression system and purified as described ^44,93,94^.

DNA fragments coding for eIF4F (6xHis-eIF4G1, eIF4A, eIF4E), 6xHis-eIF4G1, 6xHis-eIF4G2 and its C-terminally truncated form 6xHis-p86 (1-792) were obtained from human cDNA and cloned into pACEBac2 vector for MultiBac Expression system. The expression followed the previously described protocol ^94^. The cells expressing 6xHis-eIF4G1 or eIF4F complex were harvested from 800 ml of liquid culture 24 h after the day of proliferation arrest (DPA). The cells expressing 6xHis-eIF4G2 or 6xHis-tag-p86 were harvested from 400 ml of liquid culture 72 h after DPA. Cells were lysed by sonication in 20 mM Tris–HCl pH 7.5, 500 mM KCl, 1 mM DTT, 0.1% Tween-20 and 10% glycerol supplemented with protease inhibitors (Thermo Fisher Scientific, Waltham, USA). eIF4F, eIF4G1, eIF4G2 were purified by affinity chromatography using a HisTrap HP column (GE Healthcare, Chicago, USA) followed by anion-exchange chromatography using a MonoQ column in 100-500 mM gradient of KCl (GE Healthcare, Chicago, USA). eIF4E was obtained from the eIF4F preparation also using a MonoQ column in 100-500 mM gradient of KCl. p86 was purified by affinity chromatography using a HisTrap HP column (GE Healthcare, Chicago, USA) followed by anion-exchange chromatography using a MonoS column in 100-500 mM gradient of KCl (GE Healthcare, Chicago, USA).

DNA fragments coding for truncated variants of the human eIF4G – p100 (aa. 701-1600), MA3 domain (aa. 1234-1437), and W2 domain (aa. 1437-1600) were cloned to pGEX vector by BamHI / XhoI sites. All proteins were prodused in *E. coli* strain Rosetta and subsequently purified via Glutathione-Sepharose with elution by PreScission protease (Cytiva, Marlborough, USA), followed by ion-exchange chromatography on HiTrap Q (GE Healthcare, Chicago, USA).

### *In vitro* transcription of mRNA

Nluc mRNA was *in vitro* transcribed using the T7 RiboMAX Express Large Scale RNA Production System (Promega, Madison, USA) from the Nluc PCR template as described ^65^ and capped using ARCA (NEB, S1411) as described ^47^. The MVHL mRNA was transcribed *in vitro* using T7 RNA polymerase from the plasmid, which contained a T7 promoter, four CAA repeats, the β-globin 5’ UTR, a short open reading frame coding for the peptide MVHL, followed by the UAA stop codon and a 3’ UTR comprising the rest of the natural β-globin coding sequence. For run-off transcription, the MVHL-stop plasmid was linearized with XhoI endonuclease.

### Peptide release assay with NLuc

Peptide release with NLuc was performed as described previously ^67^ with minor modifications. To obtain preterminaton complexes (preTC) at the Nluc mRNA, a reaction mixture containing 70% nuclease-treated RRL (Green Hectares, Oregon, USA) was supplemented with 20 mM Hepes-KOH (pH 7.5), 80 mM KOAc, 0.5 mM Mg(OAc)_2_, 0.36 mM ATP, 0.2 mM GTP, 0.05 mM each of 20 amino acids (Promega, Madison, USA), 0.5 mM spermidine, 5 ng/µl total rabbit tRNA, 10 mM creatine phosphate, 0.003 U/µl creatine kinase (Sigma-Aldrich, Burlington, USA), 1 mM DTT and 0.2 U/µl RiboLock (Thermo Fisher Scientific, Waltham, MA USA) in a total volume of 200 µl. The mixture was preincubated with 2 μM eRF1(AGQ) at 30 °C for 10 min, followed by the addition of the NLuc mRNA to a final concentration of 8 µg/ml. Next, the mixture was incubated for 1 hour to obtain the TCs with translated Nluc, and, after that, KOAc concentration was adjusted to 300 mM and the mixture was layered on 10–35% linear sucrose gradient in a buffer containing 50 mM Hepes-KOH, pH 7.5, 7.5 mM Mg(OAc)_2_, 300 mM KOAc, and 1 mM DTT. The gradients were centrifuged in a SW55 Ti (Beckman Coulter) rotor at 55,000 rpm for 65 min. Fractions enriched with preTCs were collected by optical density and peptide release activity in the presence of excessed amounts of eRFs. PreTC was aliquoted, flash-frozen in liquid nitrogen, and stored at −80 °C.

The peptide release assay was performed in a solution containing preTC as described ^47^, 20 mM Tris-HCL, pH 7.5, 0.25 mM spermidine, 1 mM DTT, 0.2 mM GTP, and 1% Nluc substrate (Nano-Glo, Promega, Madison, USA) in the presence of release factors (50 nM of eRF1 alone, 5 nM eRF1 with 5 nM eRF3a, or 2.5 nM eRF1 with 2.5 nM eRF3c, as indicated) and 2.5-100 nM protein of interest (eIF4F, eIF4G1, eIF4G2, eIF4A, eIF4E, eIF4B, p86, p100, p50 and MA3 or W2 domains) or GST as the negative control. Luminescence was measured at 30 °C using a Tecan Infinite 200 Pro (Tecan, Männedorf, Switzerland).

### Analysis of formation of termination complexes by fluorescent toe-printing assay

PreTCs on MVHL were assembled as described previously ^56,69^. Conformation rearrangements during TC formation were detected by a fluorescent toe-print assay as described ^68,69^ with modifications. eIF4F, eIF4G1, or eIF4G2 (final concentration 125 nM), combined with eRF1(AGQ) and eRF3a (final concentrations 3 nM each) were added to preTC assembled on MVHL mRNA (final volume 10 μl) with 0.2 mM GTP. The mixture was incubated for 15 min at 37 °C. Then 2 μl of a buffer (containing 2.5 mM of each dNTPs, 25 mM Tris-HCl pH 7.5, 50 mM KAc, 2 mM DDT, and 42.5 mM MgCl_2_), 250 nM 5’-6-FAM labeled primer 5’-GCATGTGCAGAGGACAGG-3’ (Syntol, Moscow, Russian Federation), 0.625 U AMV reverse transcriptase (NEB, USA) were added. Reverse transcription was performed for 40 min at 37 °C, Then FAM-labeled cDNA products were purified by phenol-chloroform extraction. Obtained cDNAs were separated by capillary electrophoresis using standard GeneScan^®^ conditions on an Applied Biosystems® Sanger Sequencing 3500xL Genetic Analyzers (Thermo Fisher Scientific, Waltham, USA).

### GTPase assay

The 40S and 60S ribosomal subunits (250 nM each) were associated at 37 °C for 10 min, then added to a GTPase buffer (containing 10 мМ Tris-HCl pH 8.0, 12 mM NH_4_Cl, 30mM KCl, and 6 mM MgCl_2_) and 2 µM [γ – ^32^P]GTP with specific radioactive activity 2.16 μCu/pmol, to the final volume of 12 µl. Then 166 nM eRF1 were mixed with 78-312 nM of eIF4F/p50/eIF4G2. The reaction was started by addition a GTPase (166 nM eRF3a), followed by incubation at 37 °C for 10 min. Then 500 µl of 5% charcoal in 50mM Na_2_HPO_4_ were added to stop the reaction. 380 µl of liquid fraction after centrifugation at 12,000 g were applied in Perkin-Elmer scintillator counter.

### Ribosomal complex binding assay

Purified preTCs with NLuc or MVHL, as described ^61^, were incubated with 10 pmol of eRF3a, 10 pmol of eRF1(AGQ), and 10 pmol of p100 or eIF4A in a buffer containing 20 mM Tris-HCl pH 7.0, 10 mM MgCl_2_, 1 mM DTT, 0.1 mM spermidine, and 0.2 mM GTP equilibrated with 0.2 mM MgCl_2_ at 37 °C for 15 min in the volume of 100 μl. eIF4A was additionally incubated with 0.2 mM ATP, AMPNP, or without a nucleotide. The obtained complexes were centrifuged through a 10-30% (w/w) linear SDG. The whole gradient was fractionated into 14 equal fractions (385 μl), followed by precipitation with 10% (v/v) trichloroacetic acid in the presence of 2% (v/v) of casamino acids at 4 °C overnight. The protein pellets were dried, dissolved in formamide and analyzed using western blotting analysis with corresponding antibodies raised against eIF4G1 (Cell Signaling, Danvers, USA, 2858S), eIF4A (Cell Signaling, Danvers, USA, 2013S), eRF1 (Abcam, Cambridge, UK, 153731), eRF3 (Cell Signaling, Danvers, USA, 14980S), or RLP9 (Abcam, Cambridge, UK, 182556).

### Pull-down assay

HisSUMO-tagged protein (eRF3a or eRF1) or His-tagged eIF4G1, eIF4A, and eIF4B were preincubated for 15 min at room temperature with Ni-NTA Sepharose (GE Healthcare, Chicago, USA) pre-equilibrated in Wash buffer (WB, 25 mM Tris-HCl, 150 mM KCl, 10 mM imidazole, 1 mM) at a ratio of 20 pmol of protein per 1 µl of packed resin. HisSUMO or His-Acetate Kinase (Ack) bound to resin were used as a negative control. After that, this resin was washed once with WB and added 1 µl to 20 pmol of the tested protein in a buffer containing 25 mM Tris-HCl pH 7.5, 150 mM KCl, 2.5 mM MgCl_2_, 2 mM DTT, 0.25 mM spermidine, 0.2 mM nucleotide (GDP/GTP/GDPCP) supplemented with MgCl_2_. The final volume was 20 μl. The mixture was incubated for 10 min at 37 °C with constant shaking. After that, the resin was washed thrice with 500 μl of WB. The washed resin was incubated in WB with either 0.2 pmol Ulp1 protease (for His-tag removal from HisSUMO-eRF1 or HisSumo-eRF3a) in a volume of 20 µl, or 300 mM Imidazole (for elution from Ni-NTA bound with His-eIF4G1, His-eIF4A, His-eIF4B, His-Ack). After centrifugation, the supernatant was taken for electrophoretic analysis in PAAG, followed by Coomassie G-250 protein staining.

### Quantification and statistical analysis

All experiments were carried out on at least three technical replicates (the exact number of replicates is indicated in the legends). The data are presented as mean±standard deviation (SD), when analyzing luminescence signals, or mean±standard error of mean (SE), when analyzing parameters (translation). A two-tailed t-test was used to compare mean values between two groups. The Holm–Bonferroni method was used to counteract the problem of multiple comparisons.

