## Supplemental figure legends for "Eukaryotic initiation factors eIF4F and eIF4B promote translation termination upon closed-loop formation"

**SUPPLEMENTARY FIGURE LEGENDS**

**Figure S1.** (**A**) Co-expression of three human eIF4F subunits (eIF4G, eIF4A, and eIF4E) in a baculovirus expression system. The left panel shows eIF4F subunits separated by SDS-PAGE and stained by Coomassie G250. The right panel: Western blot analysis of eIF4F. The recombinant proteins expressed and isolated from baculovirus expression system Sf9 cells (eIF4G1, eIF4E) or from *E. coli* (eIF4A, eRF1, eRF3a) were used as controls (ctrl). Antibodies raised against human eIF4G1, eIF4A, eIF4E, eRF1, and eRF3 were used. (**B**) The example of the luminescence curves showing the Nluc release induced by eRF1 and eRF3a (5 nM each) in the absence and presence of eIF4F (2 nM) (the upper panel). The example of the luminescence curves shows the Nluc release induced by eRF1 (7,5 nM) in the absence and presence of eIF4F (2 nM) (the lower panel). (**C**) The influence of all studied proteins on Nluc preTC in the absence of release factors. RLU, relative luminescent units. (**D**) The peptide release rate at Nluc preTC induced by eRF1 and eRF3 (5 nM each) in the presence of various concentrations of GST as a negative control protein.

**Figure S2.** Activity of recombinant eIF4F and its subunits during translation initiation. Toe-printing analysis of 48S initiation complexes, assembled in a reconstituted translation system at capped-MVHL-polyA mRNA and uncapped-MVHL mRNA in the presence of recombinant human eIF4F, expressed and purified as multi-subunit complex in Baculovirus expression system, or eIF4F, reconstructed from individual recombinant subunits eIF4G1, eIF4A and eIF4E. Additionally 48S complexes assembly in the absence of eIF4E, eIF4A or eIF4B is shown. rfu – relative fluorescence units, nt – nucleotides.

**Figure S3.** (**A**) The luminescence curves show the Nluc release induced by eRF1 and eRF3a (5 nM each) in the absence and presence of eIF4G1, p100, and p50 (50 nM each) (1:10). (**B**) The rate of peptide release (v_0_) at Nluc preTC, induced by eRF1 (50 nM), in the presence of various concentrations of eIF4G1 (left) and difference in peptide release rates at selected concentrations: eRF1 50 nM, eIF4G 100 nM (1:2) (right). (**C**) The rate of peptide release (v_0_) at Nluc preTC induced by eRF1 and eRF3c (2.5 nM each) in the presence of at various concentrations of eIF4G1 (left) and difference in peptide release rates at selected concentrations: eRFs 2,5 nM each, eIF4G 10 nM (1:4) (right). RLU, relative luminescent units. The data are shown as the mean ± standard error, number of repeats, n = 3. Asterisks indicate statistically significant differences between the values (*, P < 0.05; **, P < 0.01; n.s., not significant).

**Figure S4.** (**A**) The luminescence curves show the Nluc release induced by eRF1 and eRF3a (5 nM each) in the absence and presence of MA3 and W2 domains (50 nM each) (1:10). (**B**) The rate of peptide release (v_0_) at Nluc preTC, induced by eRF1 (50 nM), in the presence of various concentrations of p100 (left) and difference in peptide release rates at selected concentrations: eRF1 50 nM, p100 100 nM (1:2) (right). RLU, relative luminescent units. The data are shown as the mean ± standard error, number of repeats, n = 3. Asterisks indicate statistically significant differences between the values (*, P < 0.05; **, P < 0.01; n.s., not significant).

**Figure S5.** ATPase activity of eIF4A in the absence and presence of mRNA. The data are shown as the mean ± standard error, number of repeats, n = 3. Asterisks indicate statistically significant differences between the values (**, P < 0.01).

**Figure S6.** (**A**) The luminescence curves showing the Nluc release induced by eRF1 and eRF3a (8 nM each) in the absence and presence of eIF4A or eIF4A pre-incubated with ATP (35 nM). (**B**) The luminescence curves show the Nluc release induced by eRF1 and eRF3a (8 nM each) in the absence and presence of eIF4A (35 nM) pre-incubated with AMPPNP. (**C**) The luminescence curves show the Nluc release induced by eRF1 (50 nM) in the absence and presence of eIF4A (75 nM). (**D**) The luminescence curves show the Nluc release induced by eRF1 and eRF3c (2,5 nM) in the absence and presence of eIF4A (10 nM). RLU, relative luminescent units. The data are shown as the mean ± standard error, number of repeats, n = 3.

**Figure S7.** (**A**) The rate of peptide release (v_0_) at Nluc preTC, induced by eRF1 and eRF3c (2.5 nM each), in the presence of various concentrations of eIF4A (left) and difference in peptide release rates at selected concentrations: eRFs 2.5 nM each, eIF4A 10 nM (1:4) (right). (**B**) The rate of peptide release (v_0_) at Nluc preTC induced by eRF1 and eRF3c (2.5 nM each) in the presence of various concentrations of eIF4B (left) and difference in peptide release rates at selected concentrations: eRFs 2.5 nM each, eIF4B 10 nM (1:4) (right). (**C**) The rate of peptide release (v_0_) at Nluc preTC induced by eRF1 (50 nM) in the presence of various concentrations of eIF4B (left) and difference in peptide release rates at selected concentrations: eRF1 50 nM, eIF4B 100 nM (1:2) (right). RLU, relative luminescent units. The data are shown as the mean ± standard error, number of repeats, n = 3. Asterisks indicate statistically significant differences between the values (*, P < 0.05; **, P < 0.01; n.s., not significant).

**Figure S8.** (**A**) The luminescence curves showing the Nluc release, induced by eRF1 and eRF3a (5 nM each) in the absence and presence of eIF4E (50 nM). (**B**) The luminescence curves showing the Nluc release, induced by eRF1 and eRF3a (5 nM each) in the absence and presence of eIF4B (50 nM). (**C**) The luminescence curves showing the Nluc release, induced by eRF1 and eRF3c (2.5 nM each) in the absence and presence of eIF4B (10 nM). (**D**) The luminescence curves showing the Nluc release, induced by eRF1 (50 nM) in the absence and presence of eIF4B (100 nM). RLU, relative luminescent units. The data are shown as the mean ± standard error, number of repeats, n = 3.

**Figure S9.** eIF4F stimulation of translation termination at capped and polyadenylated mRNAs. In this experiment, saturating concentrations of eIF4F were used, thus termination rates reached their maximum and were similar across all four mRNAs. (**A**) The luminescence curves showing the peptide release on preTC, assembled at cap-Nluc mRNA. (**B**) The luminescence curves showing the peptide release on preTC, assembled at Nluc mRNA. (**C**) The luminescence curves showing the peptide release on preTC, assembled at cap-Nluc mRNA with poly(A) tail (A50). (**D**). The luminescence curves showing the peptide release on preTC, assembled at Nluc mRNA with poly(A) tail (A50). The peptide release was induced by eRF1 and eRF3a (5 nM each) in the absence and presence of eIF4F (2 nM). (**E**) The peptide release rates in the absence and presence of eIF4F at different mRNAs. RLU, relative luminescent units. The data are shown as the mean ± standard error, number of repeats, n = 3. Asterisks indicate statistically significant differences between the values (*, P < 0.05; **, P < 0.01; n.s., not significant).

**Figure S10.** (**A**) Controls for possible oligomerization of eIF4A, eRF1, eRF3a under conditions of binding and centrifugation in the absence of ribosomes. Western blot analysis of the factor’s distribution in SDG. (**B**) Western blot analysis of the binding of eIF4F and release factors with the preTC. PreTCs were incubated with eRF1 or eRF3a separately or together in the presence of GTP and GDPCP (the left panel). PreTCs were incubated with p100; eRF1 or eRF3a separately or together in the presence of GTP and GDPCP (the right panel). The fractions corresponding to preTC are in a green frame. Antibodies raised against eRF1, eRF3a, RPL9, and p100 were used.

**Figure S11.** Binding of release factors with eIFs in solution. (**A**) Pull-down assay using His-tagged eIF4A and eIF4B, immobilized on Ni-NTA Sepharose and incubated with eRF1 and eRF3a separately or together. The eluted probes were analyzed by Western blot with appropriate antibodies. (**B**) Pull-down assay using HisSUMO-tagged protein (eRF3a or eRF1), immobilized on Ni-NTA sepharose and incubated with eIF4E. The washed and eluted probes were analyzed by SDS-PAGE and Coomassie G250 staining. (**C**) Pull down assay using His-tagged eIF4G, immobilized on Ni-NTA sepharose and incubated with eRF1 and eRF3a separately or together. The eluted probes were analyzed by Western blot with appropriate antibodies. Antibodies raised against eIF4G, eIF4E, eIF4A, eIF4B, eRF1, eRF3, and HIS-tag were used. GTP was added to each reaction mixture. Ctr - HisSUMO or His-Acetate Kinase (Ack) bound to resin was used as a negative control.

**Figure S12.** (**A**) Pull-down assay using HisSUMO-eRF3a bound to the Ni-NTA beads and the proteins of interest: eRF1 and p50, in the presence of GTP, GDPCP, and GDP. (**B**) Pull-down assay using HisSUMO-eRF3a bound to the Ni-NTA beads and the proteins of interest: eRF1 and p100, in the presence of GTP, GDPCP, and GDP. (**C**) Pull-down assay using HisSUMO-eRF1 bound to the Ni-NTA beads and the proteins of interest: eRF3a and MA3, in the presence of GTP, GDPCP, and GDP. The eluted probes were analyzed by SDS-PAGE and Coomassie G250 staining.
