## Supplemental figures for "Eukaryotic initiation factors eIF4F and eIF4B promote translation termination upon closed-loop formation"

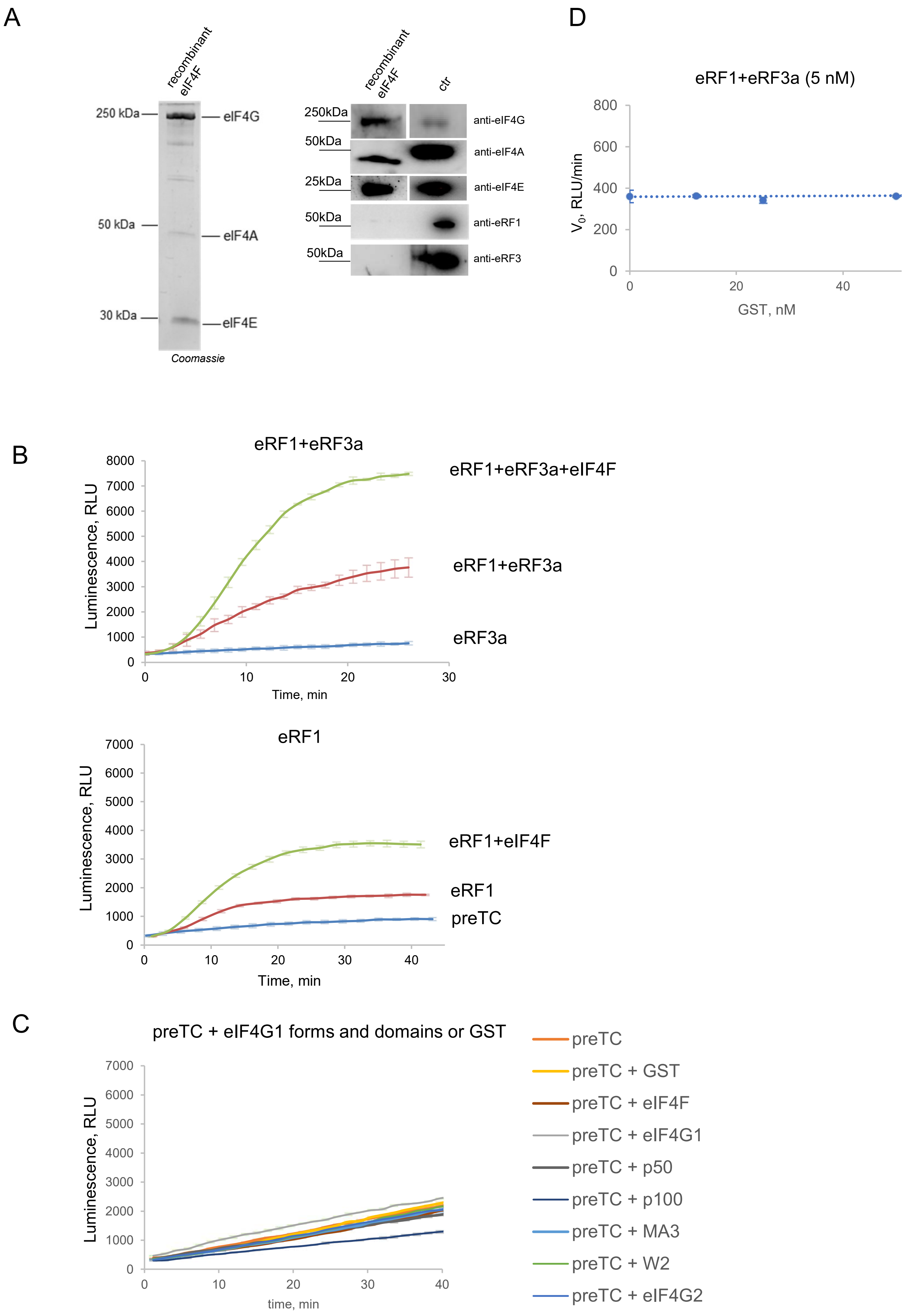
A
recombinant eIF4F
250 kDa
50 kDa
30 kDa
eIF4G
eIF4A
eIF4E
Coomassie
recombinant eIF4F
ctr
250kDa
50kDa
25kDa
50kDa
50kDa
anti-eIF4G
anti-eIF4A
anti-eIF4E
anti-eRF1
anti-eRF3
D
eRF1+eRF3a (5 nM)
V0, RLU/min
800
600
400
200
0
0
20
40
GST, nM
B
eRF1+eRF3a
Luminescence, RLU
8000
7000
6000
5000
4000
3000
2000
1000
0
0
10
20
30
Time, min
eRF1+eRF3a+eIF4F
eRF1+eRF3a
eRF3a
eRF1
Luminescence, RLU
7000
6000
5000
4000
3000
2000
1000
0
0
10
20
30
40
Time, min
eRF1+eIF4F
eRF1
preTC
C
preTC + eIF4G1 forms and domains or GST
Luminescence, RLU
7000
6000
5000
4000
3000
2000
1000
0
0
10
20
30
40
time, min
preTC
preTC + GST
preTC + eIF4F
preTC + eIF4G1
preTC + p50
preTC + p100
preTC + MA3
preTC + W2
preTC + eIF4G2

Fig. S1

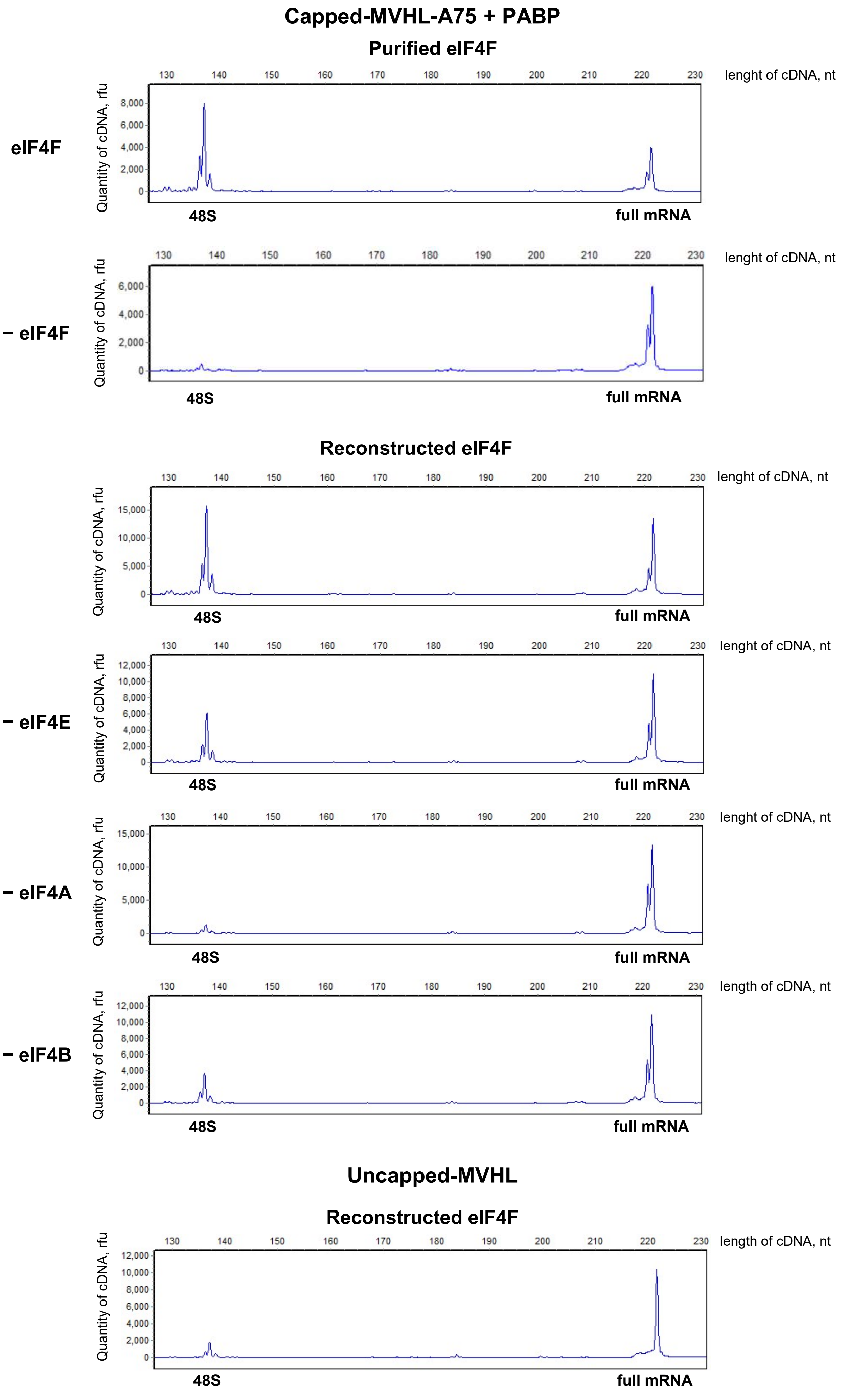

Capped-MVHL-A75 + PABP
Purified eIF4F
eIF4F
− eIF4F
Reconstructed eIF4F
− eIF4E
− eIF4A
− eIF4B
Uncapped-MVHL
Reconstructed eIF4F
130 140 150 160 170 180 190 200 210 220 230
lenght of cDNA, nt
length of cDNA, nt
Quantity of cDNA, rfu
0 2,000 4,000 6,000 8,000 10,000 12,000
0 5,000 10,000 15,000
48S
full mRNA


Fig. S2

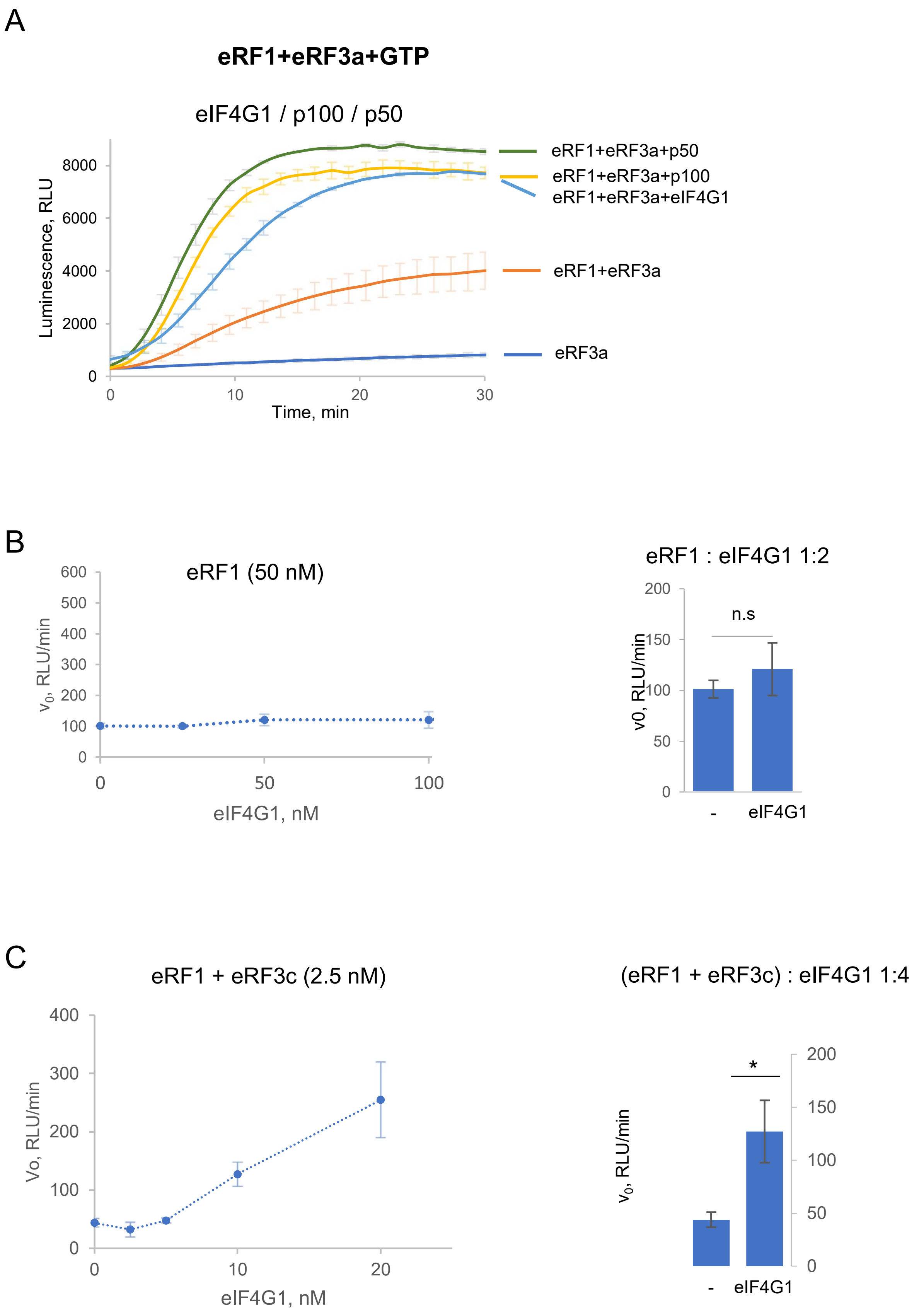
A
eRF1+eRF3a+GTP
eIF4G1 / p100 / p50
Luminescence, RLU
8000
6000
4000
2000
0
0
10
20
30
Time, min
eRF1+eRF3a+p50
eRF1+eRF3a+p100
eRF1+eRF3a+eIF4G1
eRF1+eRF3a
eRF3a
B
eRF1 (50 nM)
$v_0$, RLU/min
600
500
400
300
200
100
0
0
50
100
eIF4G1, nM
eRF1 : eIF4G1 1:2
v0, RLU/min
200
150
100
50
0
n.s
-
eIF4G1
C
eRF1 + eRF3c (2.5 nM)
Vo, RLU/min
400
300
200
100
0
0
10
20
eIF4G1, nM
(eRF1 + eRF3c) : eIF4G1 1:4
$v_0$, RLU/min
200
150
100
50
0
*
-
eIF4G1

Fig. S3

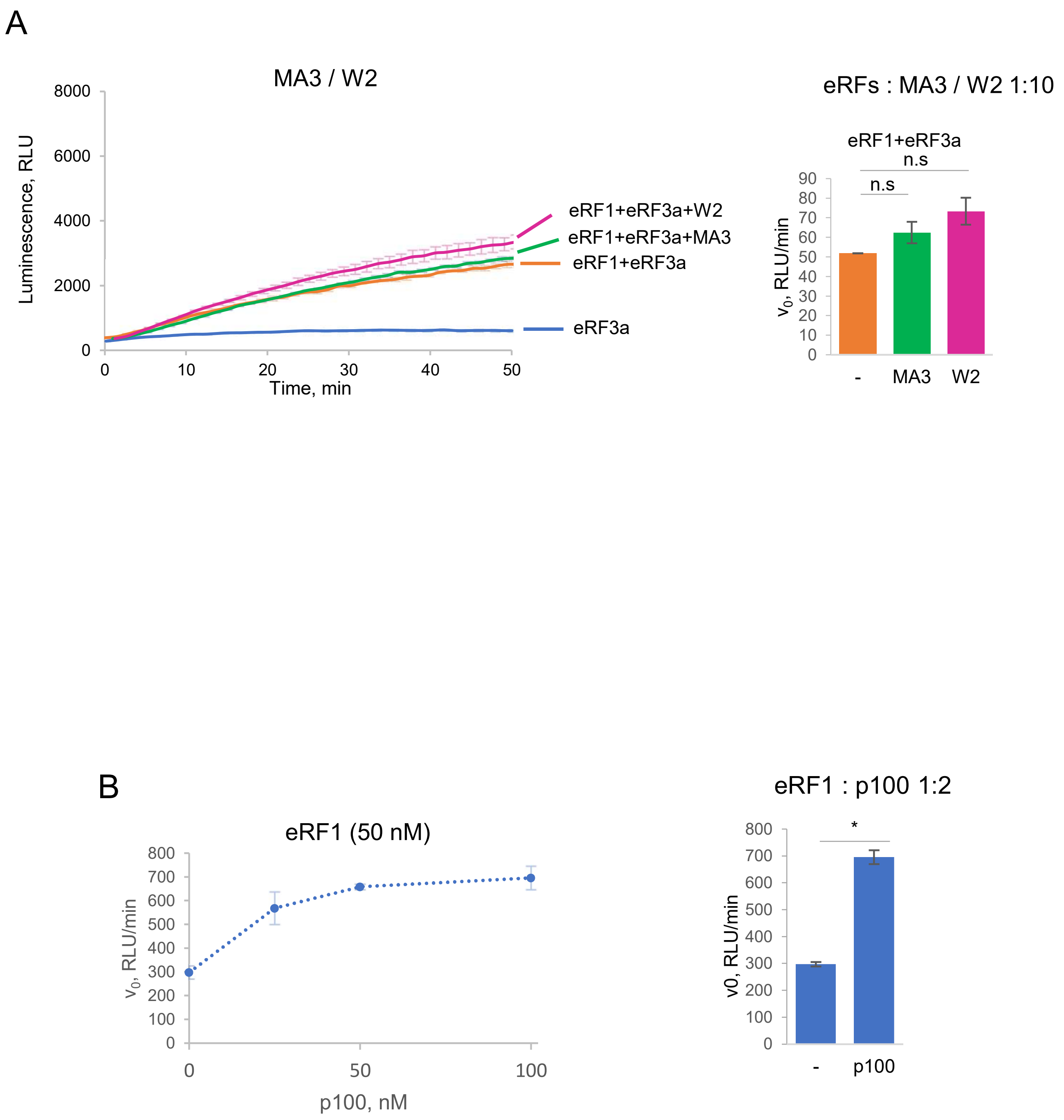
A
MA3 / W2
Luminescence, RLU
8000
6000
4000
2000
0
0
10
20
30
40
50
Time, min
eRF1+eRF3a+W2
eRF1+eRF3a+MA3
eRF1+eRF3a
eRF3a
eRFs : MA3 / W2 1:10
eRF1+eRF3a
n.s
n.s
v0, RLU/min
90
80
70
60
50
40
30
20
10
0
-
MA3
W2
B
eRF1 (50 nM)
v0, RLU/min
800
700
600
500
400
300
200
100
0
0
50
100
p100, nM
eRF1 : p100 1:2
*
v0, RLU/min
800
700
600
500
400
300
200
100
0
-
p100

Fig. S4

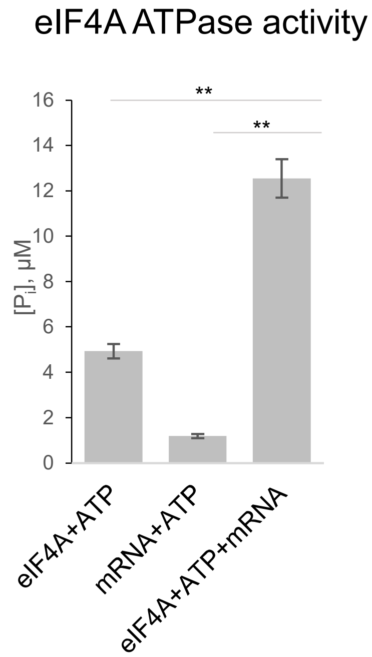
eIF4A ATPase activity
**
**
16
14
12
10
8
6
4
2
0
[Pi], μM
eIF4A+ATP
mRNA+ATP
eIF4A+ATP+mRNA

Fig. S5

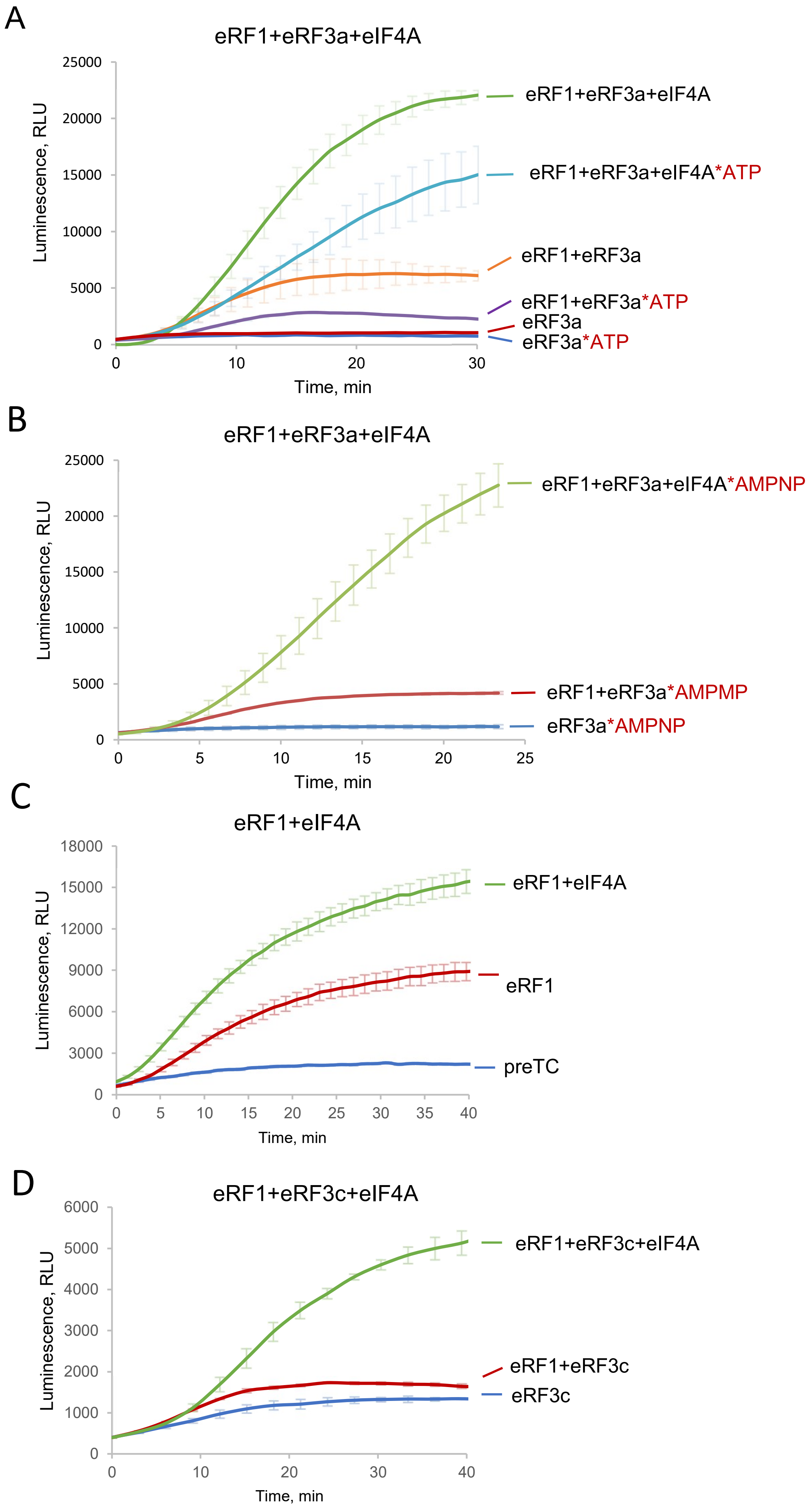
A
eRF1+eRF3a+eIF4A
Luminescence, RLU
Time, min
eRF1+eRF3a+eIF4A
eRF1+eRF3a+eIF4A*ATP
eRF1+eRF3a
eRF1+eRF3a*ATP
eRF3a
eRF3a*ATP
B
eRF1+eRF3a+eIF4A
eRF1+eRF3a+eIF4A*AMPNP
eRF1+eRF3a*AMPMP
eRF3a*AMPNP
C
eRF1+eIF4A
eRF1+eIF4A
eRF1
preTC
D
eRF1+eRF3c+eIF4A
eRF1+eRF3c+eIF4A
eRF1+eRF3c
eRF3c

Fig. S6

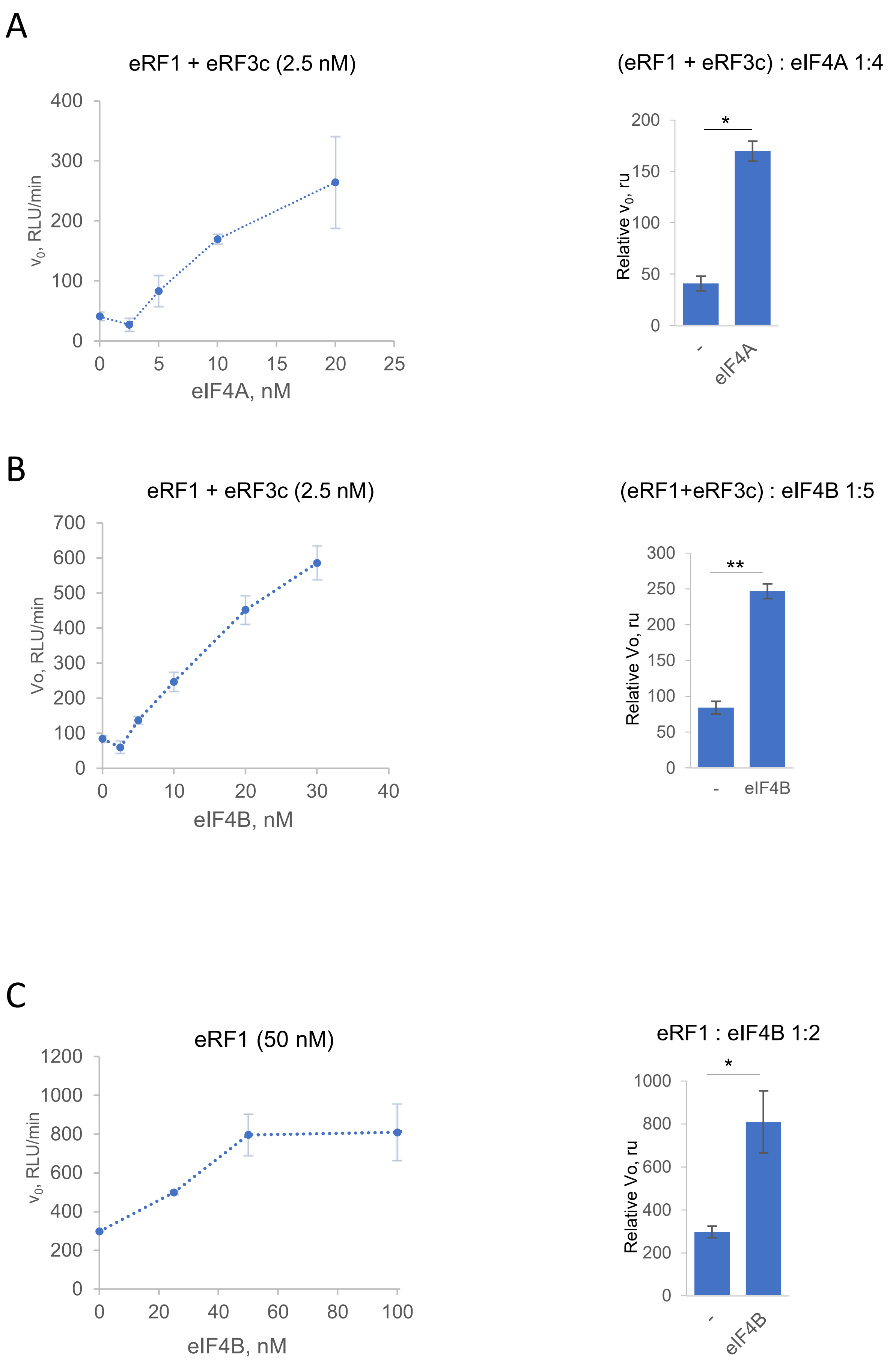
A
eRF1 + eRF3c (2.5 nM)
v0, RLU/min
400
300
200
100
0
0
5
10
15
20
25
eIF4A, nM
(eRF1 + eRF3c) : eIF4A 1:4
Relative v0, ru
200
150
100
50
0
*
-
eIF4A
B
eRF1 + eRF3c (2.5 nM)
Vo, RLU/min
700
600
500
400
300
200
100
0
0
10
20
30
40
eIF4B, nM
(eRF1+eRF3c) : eIF4B 1:5
Relative Vo, ru
300
250
200
150
100
50
0
**
-
eIF4B
C
eRF1 (50 nM)
v0, RLU/min
1200
1000
800
600
400
200
0
0
20
40
60
80
100
eIF4B, nM
eRF1 : eIF4B 1:2
Relative Vo, ru
1000
800
600
400
200
0
*
-
eIF4B

Fig. S7

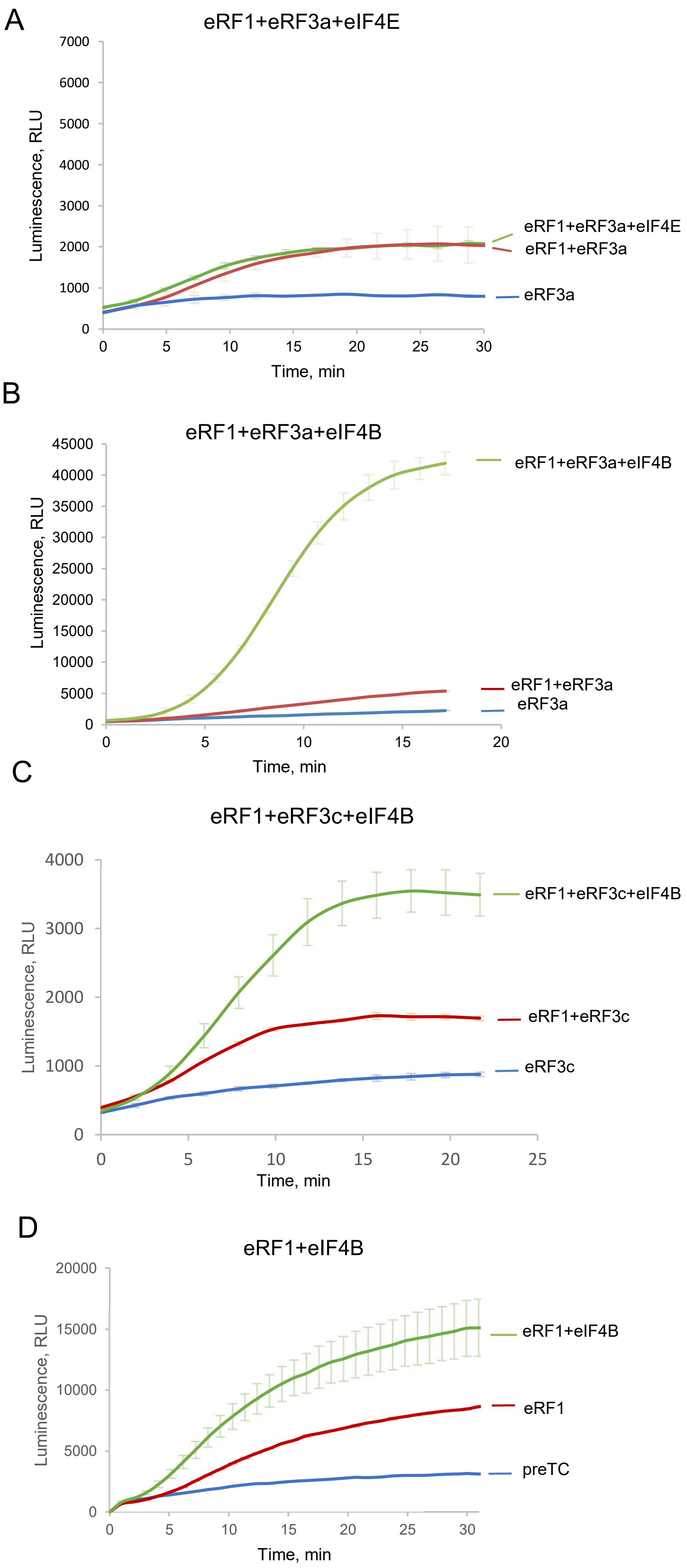
A
eRF1+eRF3a+eIF4E
Luminescence, RLU
7000
6000
5000
4000
3000
2000
1000
0
0
5
10
15
20
25
30
Time, min
eRF1+eRF3a+eIF4E
eRF1+eRF3a
eRF3a
B
eRF1+eRF3a+eIF4B
Luminescence, RLU
45000
40000
35000
30000
25000
20000
15000
10000
5000
0
0
5
10
15
20
Time, min
eRF1+eRF3a+eIF4B
eRF1+eRF3a
eRF3a
C
eRF1+eRF3c+eIF4B
Luminescence, RLU
4000
3000
2000
1000
0
0
5
10
15
20
25
Time, min
eRF1+eRF3c+eIF4B
eRF1+eRF3c
eRF3c
D
eRF1+eIF4B
Luminescence, RLU
20000
15000
10000
5000
0
0
5
10
15
20
25
30
Time, min
eRF1+eIF4B
eRF1
preTC

Fig. S8

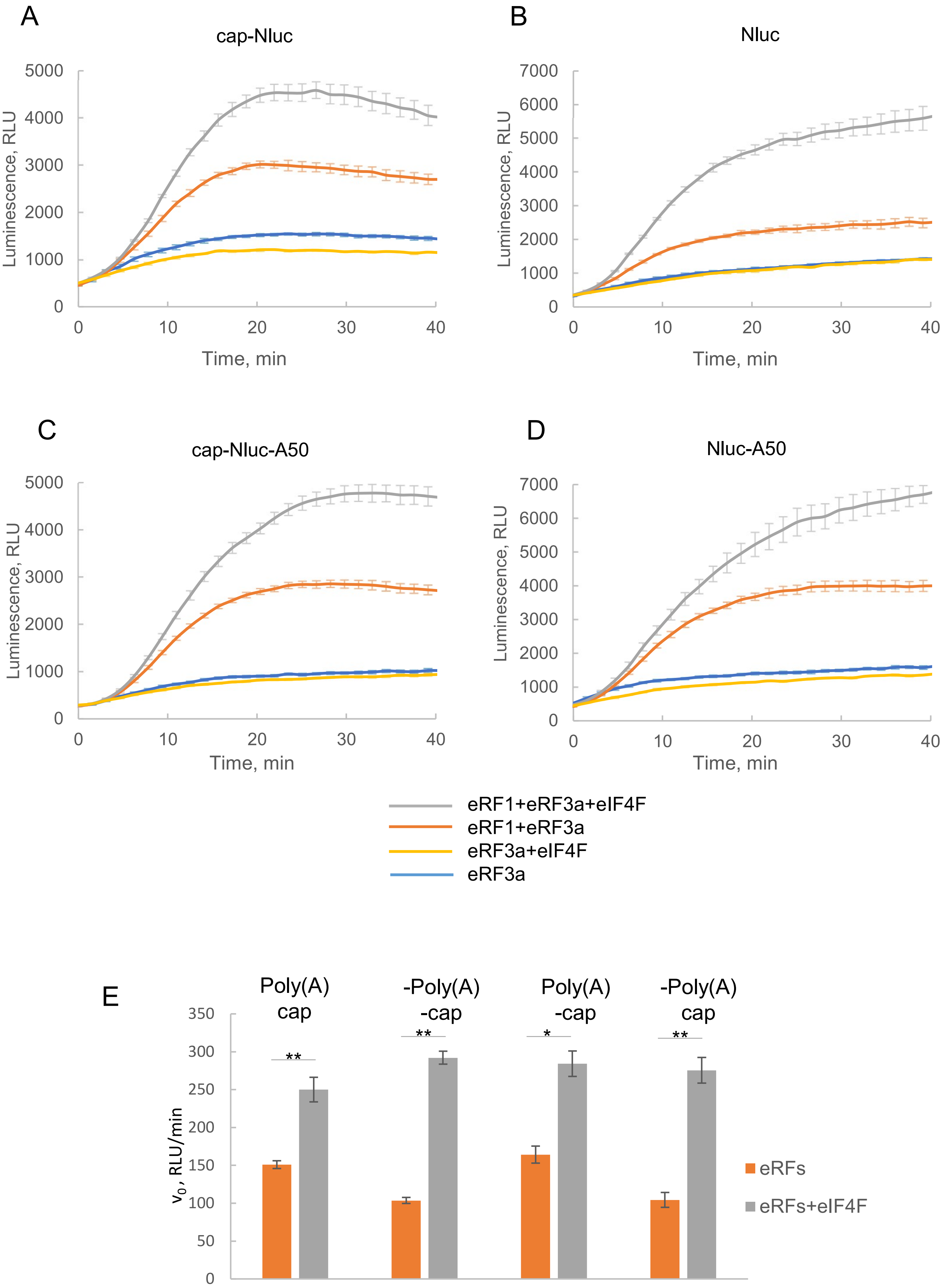
A
cap-Nluc
Luminescence, RLU
Time, min
B
Nluc
C
cap-Nluc-A50
D
Nluc-A50
eRF1+eRF3a+eIF4F
eRF1+eRF3a
eRF3a+eIF4F
eRF3a
E
Poly(A) cap
-Poly(A) -cap
Poly(A) -cap
-Poly(A) cap
v0, RLU/min
eRFs
eRFs+eIF4F

Fig. S9

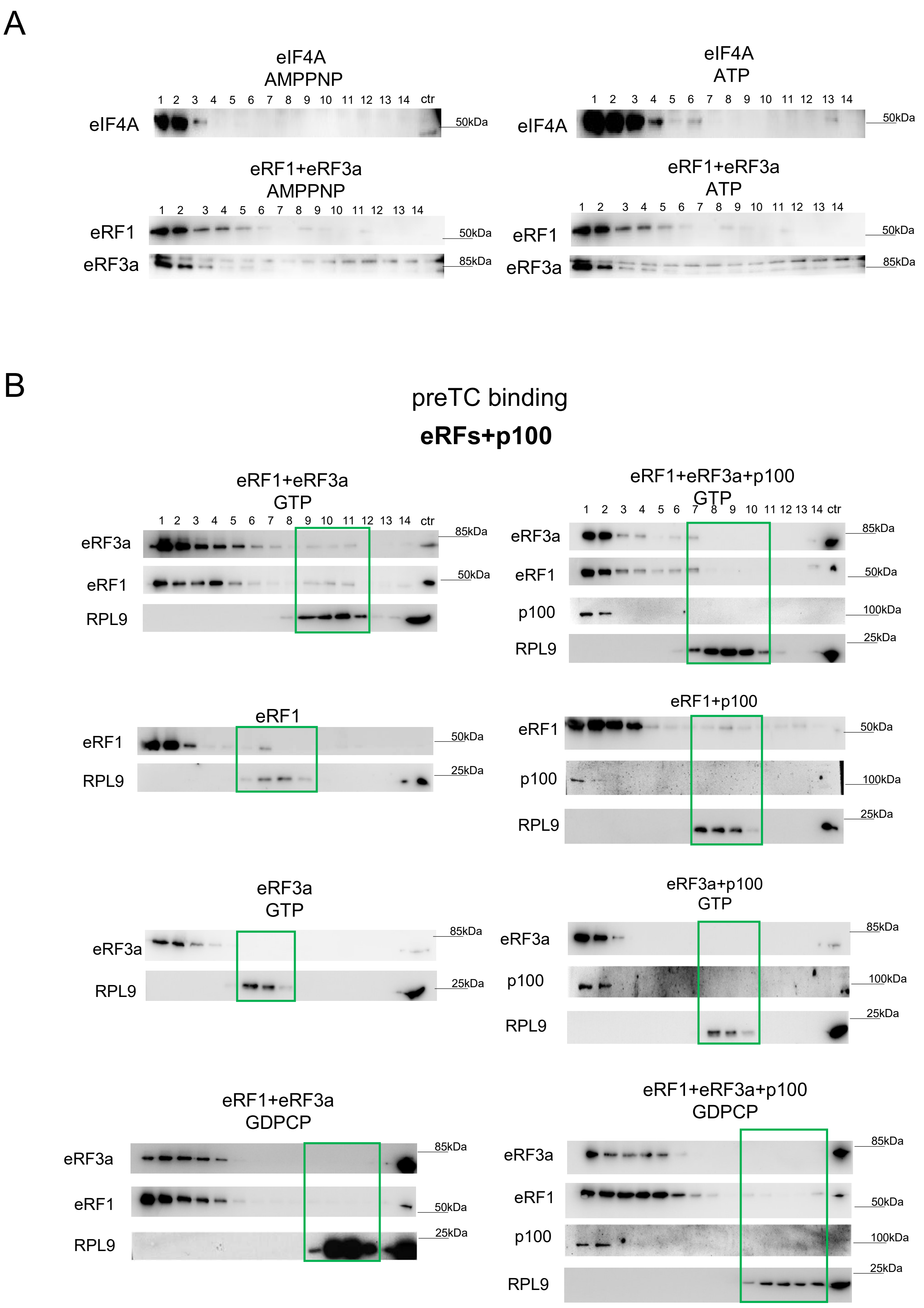
A
eIF4A
AMPPNP
1 2 3 4 5 6 7 8 9 10 11 12 13 14 ctr
eIF4A
50kDa
eIF4A
ATP
1 2 3 4 5 6 7 8 9 10 11 12 13 14
eIF4A
50kDa
eRF1+eRF3a
AMPPNP
eRF1
eRF3a
50kDa
85kDa
eRF1+eRF3a
ATP
eRF1
eRF3a
B
preTC binding
eRFs+p100
eRF1+eRF3a
GTP
eRF3a
eRF1
RPL9
85kDa
50kDa
eRF1+eRF3a+p100
GTP
p100
100kDa
25kDa
eRF1
eRF1+p100
eRF3a
GTP
eRF3a+p100
GTP
eRF1+eRF3a
GDPCP
eRF1+eRF3a+p100
GDPCP

Fig. S10

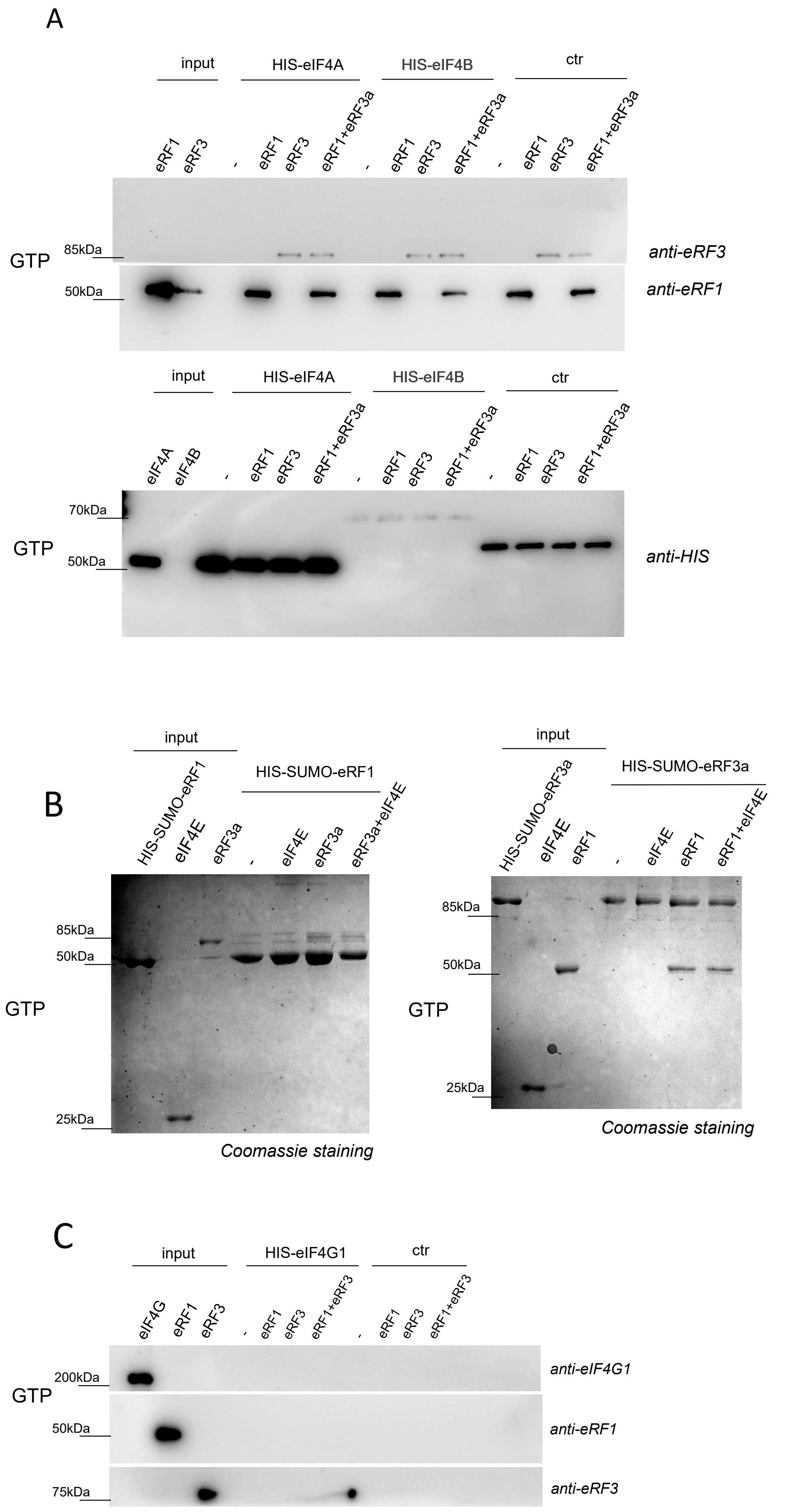
A
input
HIS-eIF4A
HIS-eIF4B
ctr
eRF1
eRF3
eRF1+eRF3a
GTP
85kDa
50kDa
anti-eRF3
anti-eRF1
eIF4A
eIF4B
70kDa
anti-HIS
B
HIS-SUMO-eRF1
eIF4E
eRF3a
eRF3a+eIF4E
25kDa
Coomassie staining
HIS-SUMO-eRF3a
eRF1+eIF4E
C
HIS-eIF4G1
eIF4G
eRF1+eRF3
200kDa
75kDa
anti-eIF4G1

Fig. S11

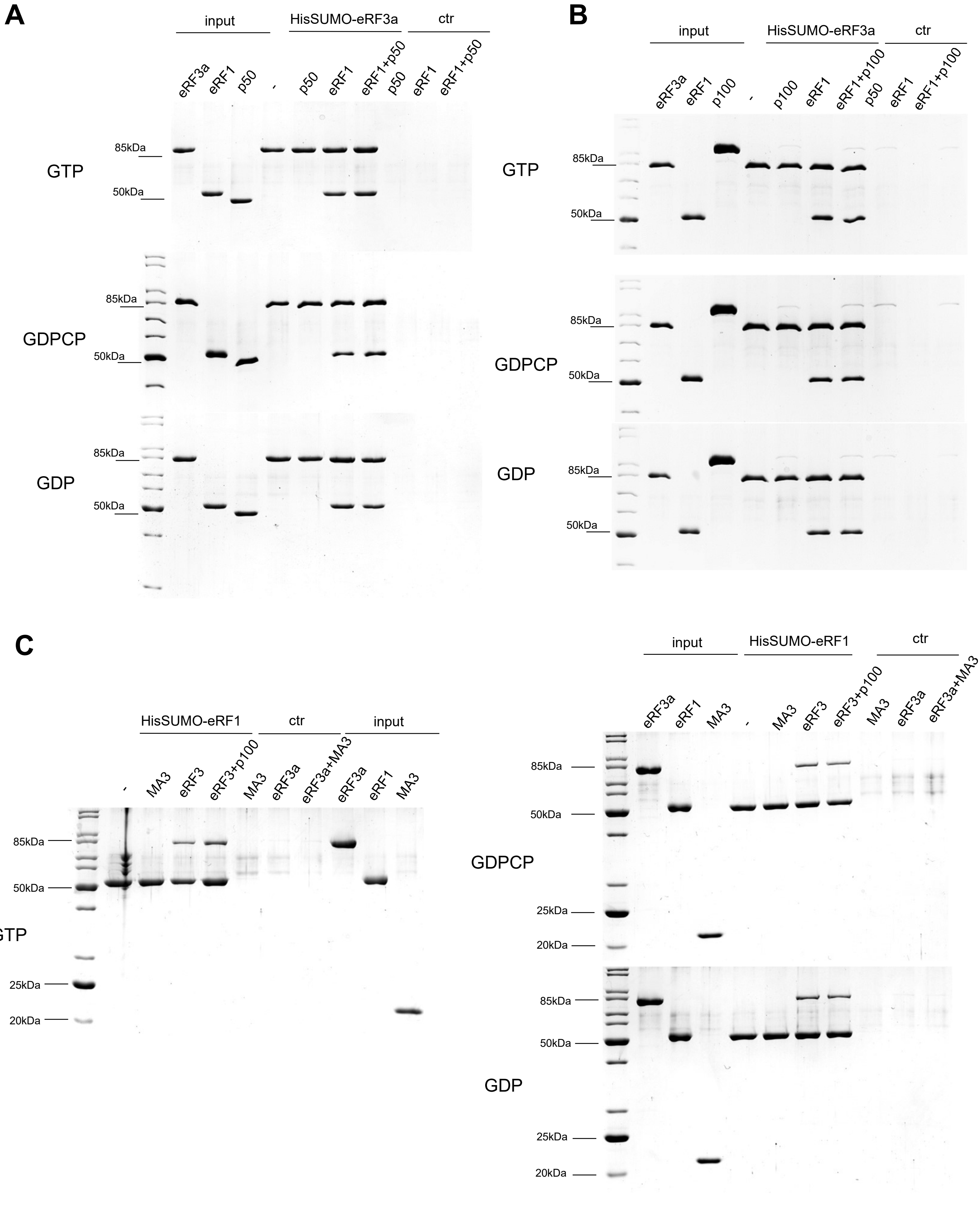
A
input
HisSUMO-eRF3a
ctr
eRF3a
eRF1
p50
-
p50
eRF1
eRF1+p50
p50
eRF1
eRF1+p50
85kDa
50kDa
GTP
GDPCP
GDP
B
input
HisSUMO-eRF3a
ctr
eRF3a
eRF1
p100
-
p100
eRF1
eRF1+p100
p50
eRF1
eRF1+p100
85kDa
50kDa
GTP
GDPCP
GDP
C
HisSUMO-eRF1
ctr
input
-
MA3
eRF3
eRF3+p100
MA3
eRF3a
eRF3a+MA3
eRF3a
eRF1
MA3
85kDa
50kDa
25kDa
20kDa
GTP
input
HisSUMO-eRF1
ctr
eRF3a
eRF1
MA3
-
MA3
eRF3
eRF3+p100
MA3
eRF3a
eRF3a+MA3
GDPCP
GDP

Fig. S12
